# Breast cancer secretes anti-ferroptotic MUFAs and depends on selenoprotein synthesis for metastasis

**DOI:** 10.1101/2023.06.13.544588

**Authors:** Tobias Ackermann, Engy Shokry, Ruhi Deshmukh, Jayanthi Anand, Laura C.A. Galbraith, Louise Mitchell, Giovanny Rodriguez-Blanco, Victor H. Villar, Britt Amber Sterken, Colin Nixon, Sara Zanivan, Karen Blyth, David Sumpton, Saverio Tardito

## Abstract

The limited availability of therapeutic options for patients with triple-negative breast cancer (TNBC) contributes to the high rate of metastatic recurrence and poor prognosis. Ferroptosis is a type of cell death caused by iron-dependent lipid peroxidation and counteracted by the antioxidant activity of the selenoprotein GPX4. Here, we show that TNBC cells secrete an anti-ferroptotic factor in the extracellular environment when cultured at high cell densities but are primed to ferroptosis when forming colonies at low density. We found that secretion of the anti-ferroptotic factors, identified as monounsaturated fatty acid (MUFA) containing lipids, and the vulnerability to ferroptosis of single cells depends on the low expression of stearyl-CoA desaturase (SCD) that is proportional to cell density. Finally, we show that the inhibition of Sec-tRNAsec biosynthesis, an essential step for selenoprotein production, causes ferroptosis and impairs the lung seeding of circulating TNBC cells that are no longer protected by the MUFA-rich environment of the primary tumour.

## Introduction

Ferroptosis is a form of cell death caused by iron-dependent peroxidation of polyunsaturated fatty acids (PUFA) within membranes. Peroxidised lipids cause stiffening and thinning of the membrane bilayer making it prone to rupture ^1,2^. In recent years, several cell-intrinsic pathways that prevent or revert lipid peroxidation have been discovered. To reduce the free radical species generated from lipidic hydroperoxide, cells rely on lipophilic antioxidants such as tetrahydrobiopterin, 7-dehydrocholesterol, ubiquinone, vitamin K and E, while the membrane-associated peroxidase selenoprotein GPX4, acts through a distinct mechanism that can directly reduce hydroperoxides ^3–10^. In addition, the fatty acid composition of the membrane influences the cellular susceptibility to ferroptosis. The synthesis of saturated fatty acids and the Acyl-CoA Synthetase Long Chain Family Member 4 (ACSL4) mediated incorporation of PUFAs into membranes promote ferroptosis ^11,12^. Furthermore, exogenous diacyl-PUFA phosphatidylcholines are pro-ferroptotic molecules that accumulate in cell membranes and correlate with cancer cell sensitivity to GPX4 inhibition ^13^. On the other hand, membrane incorporation of the exogenously supplemented or adipocyte-provided monounsaturated fatty acid (MUFA) oleic acid, as well as its cell-autonomous production mediated by the stearoyl-CoA desaturase (SCD), has been shown to prevent ferroptosis *in vitro* and *vivo* ^14–17^.

Selenocysteine is a genetically encoded amino acid, and a structural analogue of cysteine with a selenium atom instead of the sulphur one. Proteins that incorporate selenocysteine are defined as selenoproteins. During mRNA translation, selenocysteine is incorporated by a recoding event. If an mRNA contains a selenocysteine insertion sequence (SECIS), the canonical UGA stop codon is translated to selenocysteine. In addition, selenocysteine incorporation into peptides depends on the presence of specific translation factors, the eukaryotic elongation factor selenocysteine (eEFsec) and SECIS binding protein (SECISBP2)^18^. Despite the chemical similarity, cysteine and selenocysteine do not share the biosynthetic pathway. In fact, selenocysteine is synthesised on the selenocysteine tRNA (tRNA^sec^) that is firstly charged with serine by seril-tRNA synthetase. The serine on the tRNA^sec^ is then phosphorylated by phosphoseryl-tRNA kinase (PSTK). The selenophosphate (SePO (3–)) synthesized by the Selenophosphate Synthetase 2 (SEPHS2) is used by O-phosphoseryl-tRNA^Sec^ selenium transferase (SEPSECS) to synthesize selenocysteinilated tRNA^sec^ (Sec-tRNA^sec^). This is the only known pathway to produce the Sec-tRNA^sec^ and free selenocysteine obtained from selenoproteins degradation cannot be directly charged onto tRNA^sec^ ^19^.

The supplementation with an excess of selenium and the inhibition of Sec-tRNA^sec^ synthesis can both lead to the accumulation of reactive selenium species toxic to cancer cells ^20,21^. On the other hand, selenium deprivation has been shown to induce ferroptosis in cells from breast cancer, acute myeloid leukaemia, and neuroblastoma ^22–24^. Several pre-clinical and clinical studies assessed selenium supplementation as an antioxidant cancer-preventive intervention and selenium is frequently found in fortified food, and multivitamin/multimineral supplements^25,26^.

However, susceptibility to ferroptosis does not only depend on selenium availability. In this study, we show that triple-negative breast cancer (TNBC) cells under selenium starvation die of ferroptosis selectively when seeded at low density. We found that this phenotype can be rescued by transferring medium conditioned by high density cultures of breast cancer cells or cancer associated fibroblasts, but not normal cells. By means of analytical methods coupled with mass spectrometry-based lipidomics, we identified SCD-derived MUFA-containing lipids as the anti-ferroptotic factors released by TNBC cells. Finally, we interfered with the expression of genes of the selenocysteine biosynthesis pathway to prove that *in vivo* TNBC cells require the antioxidant action of selenoproteins to overcome the pro-ferroptotic environment encountered during the metastatic cascade.

## Materials and Methods

### Cell culture

Cultures of breast cancer cells (BT549, Cal120, EO771, MCF7, MDA-MB-231 and MDA-MB-468), human dermal fibroblast (DF), human mammary fibroblast (MF) and cancer associated fibroblast (CAF) were maintained in DMEM/F-12 (Thermo Fisher Scientific, #21331046) supplemented with 2 mM Glutamine (Thermo Fisher Scientific, #25030149) and 10% foetal bovine serum (FBS, Thermo Fisher Scientific, #10270106). MF and CAF cell lines were kindly provided by Prof. Akira Orimo and have been previously characterised ^27,28^. All cell lines were tested negative for mycoplasma (Venor GeM qOneStep Mycoplasma Detection Kit). Cell lines were authenticated using genomic DNA extracted with Puregene Gentra Kit and multiplexed using the Promega Geneprint Kit and multiplexed with a STR-based method (Promega Geneprint System). Samples were run on an Applied Biosystems 3130xl DNA analyser and the results analysed using the Applied Biosystems Genemapper v4.1 software. Profiles were matching the references reported by ATCC (LGC standards), Cellosaurus and DSMZ databases.

### Conditioned medium

To condition medium from high density cultures of BT549, Cal120, EO771, MCF7, MDA-MB-231, MDA-MB-468, DF, MF and CAF cell lines, 6×10^6^ cells were seeded in DMEM/F-12 with 10% FBS in a dish with a diameter of 145mm. The day after seeding, the medium was replaced with 20 ml of DMEM/F-12 without FBS and conditioned for two days. For non-targeting control (NTC) and SCDko MDA-MB-468 cells, 5×10^6^ and 1.1×10^7^ cells were seeded respectively to condition the medium. For experiments with acute SCD inhibition, 2×10^7^ MDA-MB-468 cells were seeded in DMEM/F-12 with 10% FBS in a dish with a diameter of 145mm. The day after seeding the medium was replaced by DMEM/F-12 without FBS and supplemented with 200 nM CAY10566 (SCD inhibitor, Cayman chemicals, #10012562). After 18h incubation with 200 nM CAY10566, cells were washed once with PBS, and 15 ml DMEM/F-12 medium without FBS was conditioned for 2h. For experiments shown in Figure 2A, 6×10^6^ MDA-MB468 cells were seeded in DMEM/F-12 with 10% FBS in a dish with a diameter of 145mm. The day after seeding, the medium was replaced with 20 ml of DMEM/F-12 or selenite-free Plasmax™ ^23^, both supplemented with 10% FBS and conditioned for two days. The conditioned medium was diluted with three volumes of serum-free unconditioned medium and used for the colony forming assays.

**Figure 1:**
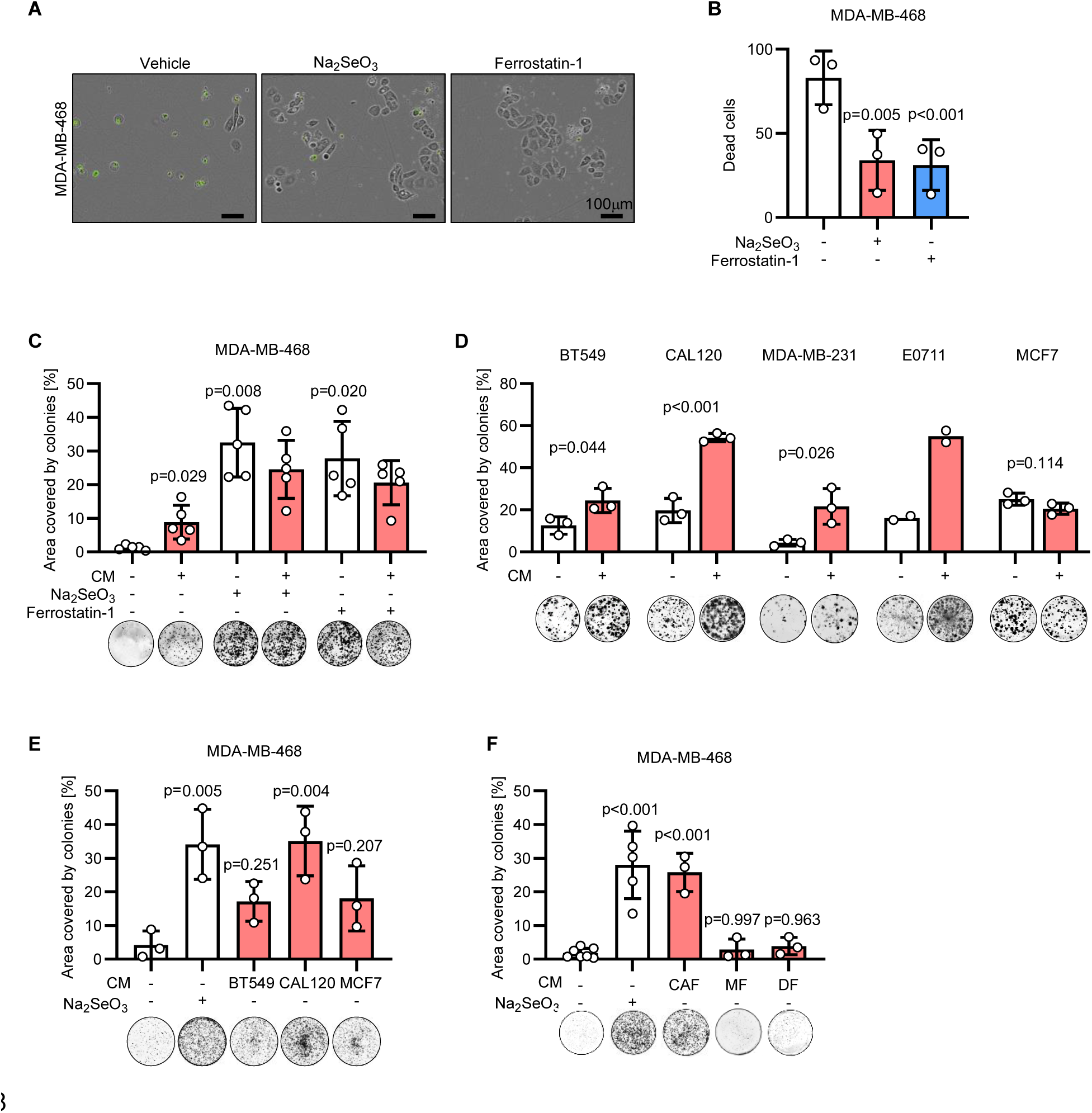
Medium conditioned by breast cancer cells and cancer-associated fibroblasts enhances the clonogenicity of triple-negative breast cancer cells. A. Representative images of MDA-MB-468 cells seeded at low density and incubated for 96 h with 50 nM Na_2_SeO_3_ or 2µM Ferrostain-1 as indicated. Phase contrast and fluorescence images were overlayed and dead cells were identified by green fluorescence emitted by the Incucyte® Cytotox Green Dye. B. Quantification of the dead cells stained with the green fluorescent dye in the conditions described in A. B. Well area covered by colonies formed by MDA-MB-468 cells incubated for 7 days with mock medium or medium conditioned by MDA-MB-468 cells seeded at high density (CM). Media used for the colony forming assay were supplemented with 50 nM Na_2_SeO_3_ or 2µM Ferrostain-1 as indicated. P values refer to a two-way ANOVA test for unpaired samples with Dunnett’s multiple comparisons test. C. Well area covered by colonies formed by the indicated cell lines incubated for 7 days with mock medium or medium conditioned by BT549, CAL-120, MDA-MB-231, EO771 and MCF7 cells seeded at high density. Conditioned medium was used on the respective cell line. P values refer to a two-tailed, homoscedastic Student’s *t* tests for unpaired samples. D. Well area covered by colonies formed by MDA-MB-468 incubated for 7 days with mock medium or medium conditioned by BT549, CAL120 or MCF7 cells seeded at high density. Media used for the colony forming assay were supplemented with 50 nM Na_2_SeO_3_ as indicated. P values refer to a one-way ANOVA test for unpaired samples with Dunnett’s multiple comparisons test. E. Well area covered by colonies formed by MDA-MB-468 incubated for 7 days with mock medium or medium conditioned by cancer associated fibroblasts (CAF), immortalised mammary fibroblasts (MF), or immortalised human dermal fibroblasts (DF) seeded at high density. Media used for the colony forming assay were supplemented with 50 nM Na_2_SeO_3_ as indicated. P values refer to a one-way ANOVA test for unpaired samples with Dunnett’s multiple comparisons test. C-F. Representative images of wells with colonies are shown for each experimental condition. n_exp_=2-7 as indicated by the data points in each panel. Bars represent mean ± s.d..

**Figure 2:**
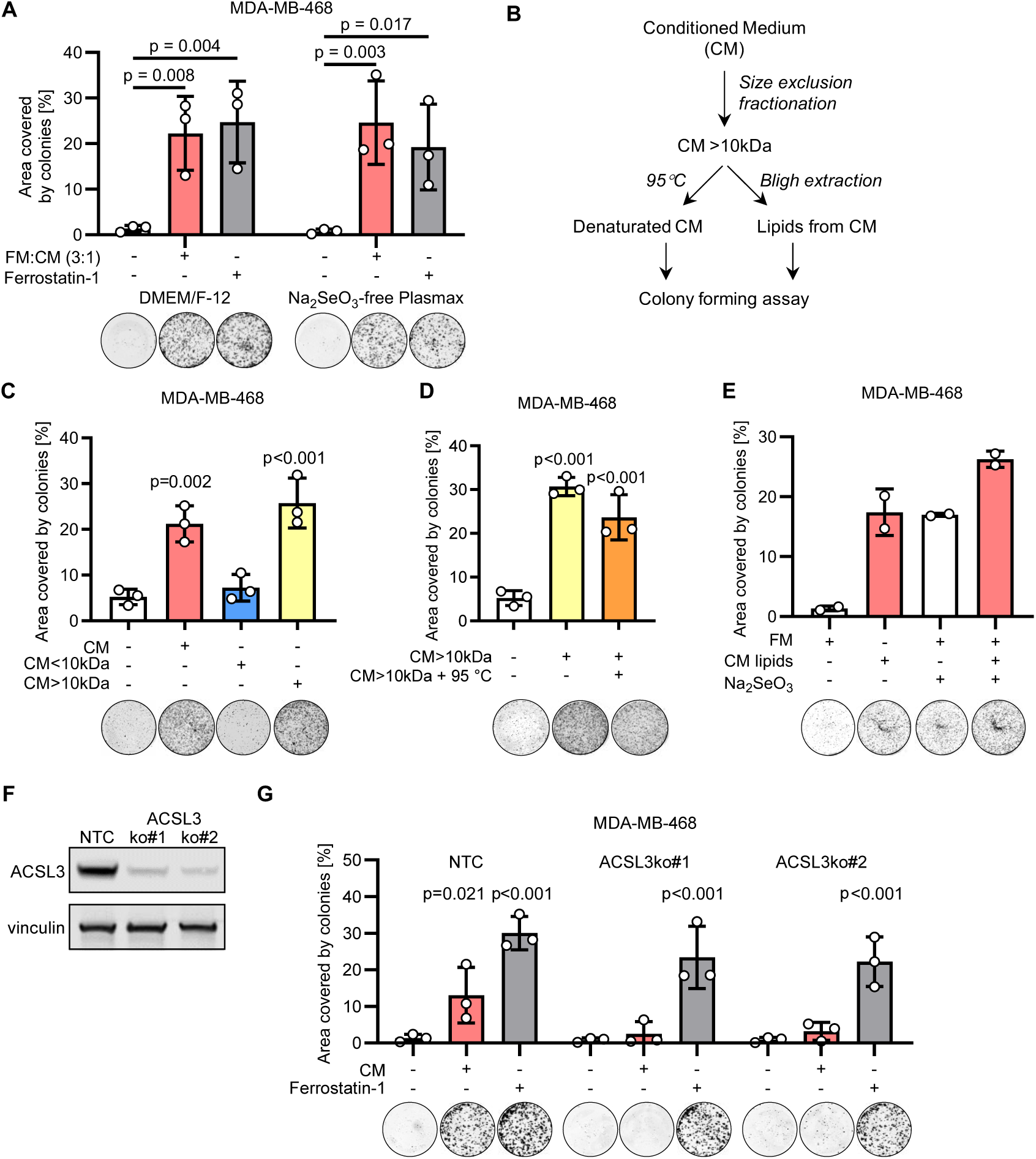
Breast cancer cells produce an anti-ferroptotic molecule at high density. A. Well area covered by colonies formed by MDA-MB-468 incubated for 7 days with fresh medium (FM) without or with supplementation of 2 µM ferrostatin-1, or with FM mixed 3:1 with medium conditioned by MDA-MB-468 cells seeded at high density (CM). Experiments were performed in DMEM/F-12 or Na_2_SO_3_-free Plasmax™. P values refer to a two-way ANOVA test for unpaired samples with Dunnett’s multiple comparisons test. B. Schematic diagram of the analytical procedures applied to CM to identify the factor rescuing colony formation. C-D. Well area covered by colonies formed by MDA-MB-468 incubated for 7 days with CM or with medium supplemented with the fractions separated by a size exclusion column as described in the Methods section (C), and further incubated for 15 min at 95°C (D). P values refer to a one-way ANOVA test for unpaired samples with Dunnett’s multiple comparisons test. E. Well area covered by colonies formed by MDA-MB-468 incubated for 7 days with fresh medium (FM) supplemented with lipidic extracts obtained from the conditioned medium (CM) as shown in B. F. Immunoblot of ACSL3 and vinculin (loading control) in NTC and two ACSL3 ko clones obtained from MDA-MB-468 cells. G. Well area covered by colonies formed by MDA-MB-468 control (NTC) or ACSL3 ko cells incubated for 7 days with fresh medium, conditioned medium (CM) or fresh medium supplemented with 2 µM ferrostatin-1 as indicated. P values refer to a two-way ANOVA test for paired samples with Sidak’s multiple comparisons test. A, C-E, G. Representative images of wells with colonies are shown for each experimental condition. n_exp_=2-3 as indicated by the data points in each panel. Bars represent mean ± s.d..

To generate mock medium, cell culture medium was incubated in a cell culture dish maintained at 37°C without cells for the respective experimental time periods (2h or 2 days).

### Colony formation assays

Colony formation assays were performed as described previously ^23^. For MDA-MB-468 cells, we seeded 5,000 cells/well in a 6 well plate with 2 ml/well of DMEM/F-12 supplemented with 2.5% FBS. To test the effects of lipid extracts from medium conditioned by MDA-MD-468 cells, 1,000 cells/well were seeded in a 24 well plate with 0.5ml of medium/ well. After 7 days incubation, colonies from MDA-MB-468 cells were fixed by replacing the medium with a solution of 3% trichloroacetic acid in water (Merck, #T6399-500G), rinsed twice with water, stained with 0.057% Sulforhodamine B solution in 1% acetic acid (Merck, #230162) and washed twice with a 1% acetic acid solution in water. For BT549, Cal120, MCF7, MDA-MB-231 and EO771 cells, 500 cells/well were seeded in a 6 well plate and fixed after 14 days of culture with the exception of EO771 cells that were fixed 7 days after seeding. Plates stained with Sulforhodamine B were scanned with Li-Cor Odyssey® DLx imaging system and the fluorescent signal quantified with ImageJ as previously described ^23^. In figures reporting colony area or number, each data point represents an independent colony forming assay mean of 2 or 3 replicate wells.

### Cell proliferation assay

To determine the number of NTC control and SCDko cells during the medium conditioning experiments, 3.32×10^5^ NTC control cells per well and 7.3×10^5^ SCDko cells per well were seeded in a 6 well plate with DMEM/F-12 containing 10% FBS. One day after seeding, cells were trypsinized and counted with a Casy counter. At this time, the medium of parallel plates seeded and incubated in the same conditions was replaced with DMEM/F-12 without FBS, and two days after, the cells were counted with a Casy counter.

For proliferation assays, NTC control or SCDko MDA-MB-468 cells were seeded at 3×10^4^/ well in a 6 well plate with DMEM/F-12 supplemented with 2.5% FBS, 50 nM Na_2_SeO_3_ (Merck, #S5261), 2.5 µM Deferoxamine (Merck, # D9533), 1 µM Ferrostatin-1 (Merck, #SML0583), 10 µM oleic acid (Merck, #O1008), or vehicles (0.05% DMSO and 0.02% ethanol). For proliferation assays with GPX4 inhibitor (RSL3), cells were seeded and incubated as described above and supplemented with 50 nM RSL3 (Merck, #SML2234) and 10% FBS. After 5 days, cells were counted with a Casy counter or stained with Sulforhodamine B and images acquired and analysed as described for the colony formation assays.

### Live cell imaging

For Live cell imaging, MDA-MB-468 cells were seeded at 5×10^3^/ well in a 24 well plate with DMEM/F-12 containing 2.5% FBS and 3 nM Incucyte^®^ Cytotox Green Dye (Satorius, #4633), 50 nM Na_2_SeO_3_ or 2 µM Ferrostatin-1 as indicated in figures. 24h after seeding and every 2h thereafter, 9 images per well were acquired with phase contrast and green fluorescence (acquisition time: 200 ms) with a 10x objective using an Incucyte S3 (Satorius). To quantify the number of dead cells, the fluorescent objects were counted using Incucyte 2022B software.

### Medium fractionation and heat inactivation

Medium conditioned by MDA-MB-468 cells was loaded into size exclusion columns (Amicon® Ultra-15 Ultracel-10 Centrifugal Filter Unit, Merck, #UFC901024) centrifuged at 4,000g for 30 min and the fractions stored at -20⁰C until further analysis. The concentrated fraction (CM>10kDa) was diluted 1:40 in unconditioned DMEM/F-12 medium, supplemented with 2.5% FBS and used for colony forming assays. The column flow-through (CM<10kDa) was supplemented with 2.5% FBS and used undiluted for colony forming assays.

For heat inactivation and protein denaturation, the concentrated fraction (CM>10kDa) was incubated for 15 min at 95⁰C, afterwards diluted 1:40 in unconditioned DMEM/F-12 medium supplemented with 2.5% FBS and used for colony forming assays.

### Lipid extraction

Lipids were extracted from the concentrated fraction (CM>10kDa) of the conditioned medium following the Bligh and Dyer method and used for colony forming assays or directly from the conditioned medium with a methyl tertiary-butyl ether (MTBE) solution and used for lipidomic analysis^29^. For the Bligh and Dyer extraction, 250 µl of the concentrated fraction (CM>10kDa) from the size exclusion columns were mixed with 960 µl of a 1:2 chloroform:methanol mixture. Afterwards, 310 µl chloroform and 310 µl water were added and the solution mixed to achieve a phase separation. The lower, chloroform containing phase was transferred to a new glass vial, dried at room temperature by nitrogen flow and resuspended in 25 µl of ethanol. 5 µl of lipid containing ethanol was supplemented to 2 ml of DMEM/F-12 with 2.5% FBS to perform colony forming assays.

For the MTBE extraction, 5 ml MTBE and 1 ml methanol were added to 2.5 ml of conditioned medium. After mixing the solution three times for 30 s, the upper phase was transferred to a new glass vial, dried under nitrogen flow at 35⁰C and resuspended in 25µl ethanol. 1 µl of the lipid ethanol solution was supplemented to 1 ml of DMEM/F-12 with 2.5% FBS and used for colony forming assays.

### Lipidomic analysis

For lipidomic analysis, 5×10^5^ NTC control and SCDko cells were seeded in 20 ml of DMEM/F-12 supplemented with 2.5% FBS in plates of 145 mm diameter. After 2 days, cells were scraped off the plate in the culture medium, collected in a tube and centrifuged at 1,000g for 3 min. The cell pellet was resuspended in 1 ml of ice-cold PBS, transferred to 1.5 ml Eppendorf vial, and centrifuged at 10,000 g for 10 s. Lipids were extracted from the cell pellet with 200 µl of butanol:methanol solution (1:1) centrifuged at 16,000g for 10 min, and analysed using high resolution mass spectrometry.

For lipidomic analysis of low-density cultures, 1.6×10^5^ MDA-MB-468 cells were seeded in plates of 145 mm diameter with 20 ml of mock or conditioned DMEM/F-12 supplemented with 2.5% FBS and 2 µM Ferrostatin-1. Vehicle control or 10 µM oleic acid were added as indicated in Figure S3A. After 2 days, cells were scraped off the plate in the culture medium, collected in a tube and centrifuged at 1,000g for 3 min. The cell pellet was resuspended in 1 ml of ice-cold PBS, quickly transferred to 1.5 ml Eppendorf vial, and centrifuged at 10,000 g for 10 s. Lipids were extracted from the cell pellet with 100 µl of butanol:methanol solution (1:1), centrifuged at 16,000g for 10 min, and analysed using high resolution mass spectrometry. For the lipidomic analysis of media, 10 µl of conditioned medium enriched (CM>10kDa) or depleted (CM<10kDa) fractions were diluted with 190 µl of butanol:methanol solution (1:1), centrifuged at 16,000g for 10 min, and analysed using high resolution mass spectrometry. For the lipidomic analysis of interstitial fluid (IF), 5µl of IF from individual NTC or SCDko tumours were diluted with 45 µl of butanol:methanol solution (1:1), centrifuged at 16,000g for 10 min, and analysed using high resolution mass spectrometry.

Lipidomic analyses were performed using a Thermo Fisher Scientific Ultimate 3000 binary UPLC coupled to a Q Exactive Orbitrap mass spectrometer equipped with a Heated Electrospray Ionization (HESI-II) source (Thermo Fisher Scientific, Massachusetts, USA). For each sample, MS data were acquired using full MS/ dd-MS2 in positive and negative modes to maximize the number of detectable species. Details on the parameters for the MS methods using different polarities is provided in Table 1. Chromatographic separation was achieved using a Waters CSH C18 analytical column (100 x 2.1mm, 1.7µm) maintained at 55°C. The mobile phase consisted of 60:40 (v/v) acetonitrile:water containing 10 mM ammonium formate and 0.1% formic acid (phase A) and 90:10 (v/v) isopropanol:acetonitrile containing 10 mM ammonium formate and 0.1% formic acid (phase B) at a flow rate of 400 µl/min. The gradient elution consisted of 30% B for 0.5 min, increasing linearly to reach 50% at 4 min, then 80% at 12 min, then 99% B at 12.1 min then held at 99% for 1 min, then returned to starting condition in 1 min and kept constant for 2 min. The total run time was 20 min. The whole system was controlled by Xcalibur version 4.3. Quality control (QC) samples prepared by mixing equal volumes of experimental samples were injected at regular interval throughout the whole batch to monitor the instrument performance. Lipidomics data were analysed with Compound Discoverer v.3.1 (ThermoFisher Scientific) and LipiDex^30^, an open-source software suite available at http://www.ncqbcs.com/resources/software/. Briefly, raw files were loaded into Compound Discoverer and processed using two workflows (aligned and unaligned) as previously described ^31^. Compound result tables were exported for further processing using the ‘Peak picking’ tab in Lipidex. In addition, the raw data files were converted to .mgf files using MSConvert (ProteoWizard, P. Mallick, Stanford University) ^32^ and imported into the ‘Spectrum Searcher’ tab in Lipidex, where the following libraries were searched ‘LipidBlast_Formate’, ‘LipiDex_HCD_Formate’, ‘LipiDex_Splash_ISTD_Formate’, ‘LipiDex_HCD_ULCFA’ using the default search tolerances for MS1 and MS2. For a lipid to be considered identified it required a minimum of 75% spectral purity, an MS2 search dot product score of at least 500 and reverse dot product of at least 700.

**Table 1.**
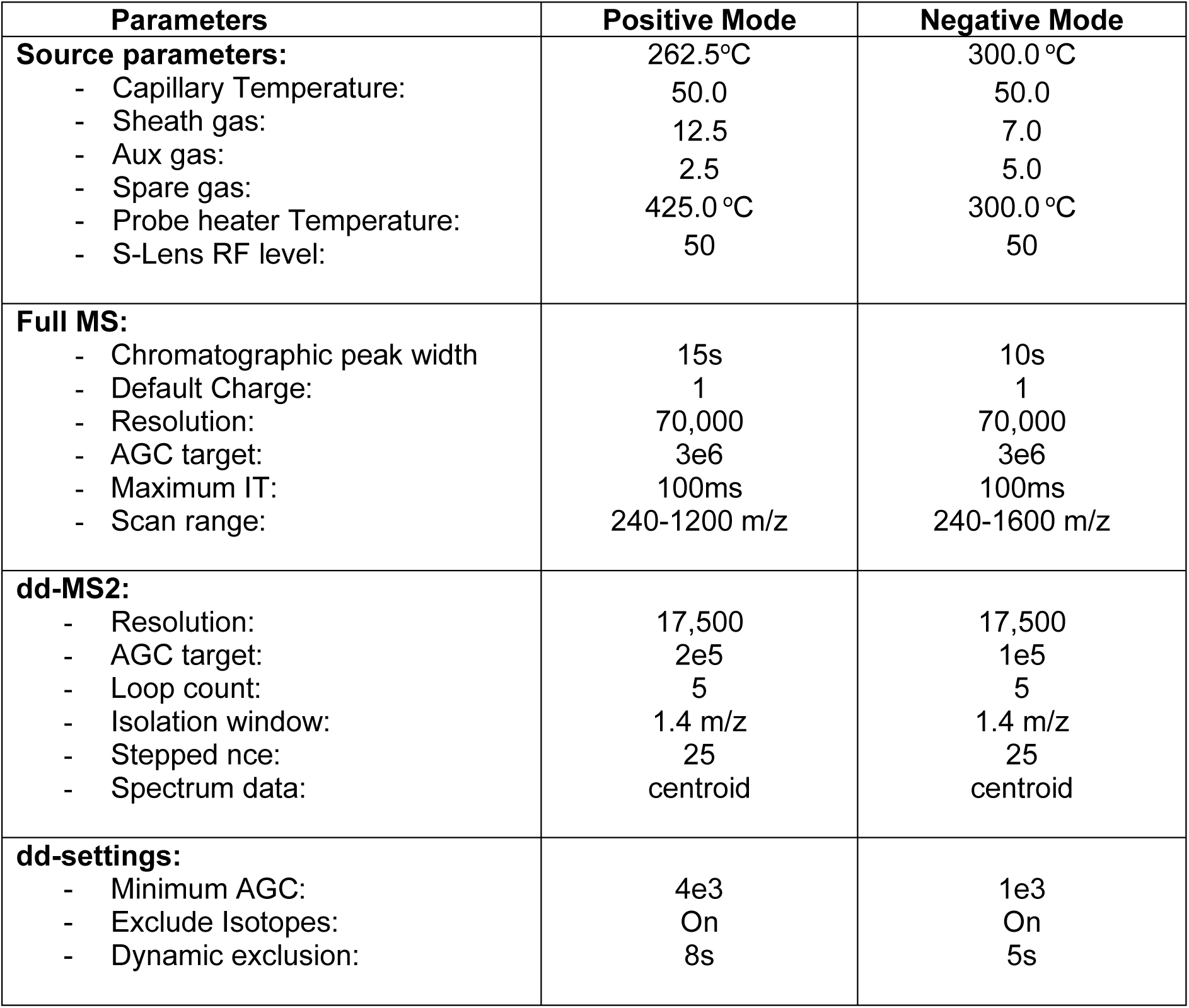
Parameters of the mass spectrometry methods using positive and negative polarities.

For the determination of the total fatty acid content of the lipid species, samples were analysed using GC-MS after transesterification using a methanolic solution of (m-trifluoromethylphenyl) trimethyl ammonium hydroxide (Meth-Prep II), to provide fatty acid methyl esters (FAME) in a one-step reaction followed by monitoring of the generated FAME using GC/MS in SIM mode. Briefly, samples were first spiked with heptadecanoic (C17:0) as an internal standard, dried under nitrogen stream and resuspended in 27µL of chloroform. 3 µL of Meth-Prep II were added and samples were mixed and analysed within 2 days. Analysis was performed using an Agilent 7890 GC chromatograph and 7693 autosampler coupled with an Agilent 7000 GC/MS provided with (Agilent, CA, United states). High purity helium (99.999%) was used as carrier gas at an initial flow rate of 1 mL/min increased to 2 mL/min in 12 seconds and kept constant for 1 min and then returned to 1mL/min for the rest of the run time. The chromatographic column used was Phenomen BPX70 (60 m x 250 µm, 0.25 µm) (Fisher Scientific, USA). The injector was set at a temperature of 300 °C, a pressure of 21 psi and a septum purge flow of 3mL/min. Injection was performed in a pulsed splitless mode at 60 psi for 1.2 min and a purge flow to split vent of 50 mL/min at 1.2 min. The injection volume was 1 µl. The oven temperature was set to an initial temperature of 100°C held for 2 min followed by a linear increase to 172°C at 8°C/min (held for 6 min), then 196°C (held for 9 min) then 204°C (held for 15 min). The total run time was 45 min. For the separation of the palmitoleic acid and oleic acid isomers, a longer chromatographic run was used applying the same starting conditions, followed by a stepwise increase in temperature initially to 160°C at 8°C/min (held for 6 min), then to 185 °C at 0.5°C/min (held for 9 min), then 204°C at 8°C/min (held for 15 min). The MS was operated in electron ionization (EI) mode at 70 eV and the MSD transfer line was set to 260 °C. MS data were acquired in SIM mode for monitoring of the individual fatty acid derivatives. MS recording started at a cut-off of 8 min. Data were processes using Agilent MassHunter Quantitative analysis software (Agilent, CA, United states). Initially, fatty acids were annotated by comparison to FAME Standard Mixture (Merck Life Science UK Limited, United Kingdom) using a combination of retention time and fragmentation pattern using full scan. A SIM mode for selected fragments was applied to improve the sensitivity of detection.

### Cloning and CRISPR-based gene editing

The following gRNA sequences: Non-Targeting Control (NTC): 5’-GTAGCGAACGTGTCCGGCGT-3’, ACSL3 5’-GAGCTATCATCCACTCGGCCC-3’, LRP8 5’-GGCCACTGCATCCACGAACGG-3’, *PSTK*: 5’-AAACTGATCAGACACTCCGA-3’, *SCD*: 5’-GCAGCCGAGCTTTGTAAGAG-3’, *SEPHS2*: 5’-GAGGGACGGCAGTGACCGG-3’, *SEPSECS*: 5’-AACCGCGAGAGCTTCGCGG-3’ were cloned into lentiCRISPRv2 vector (Addgene, Plasmid #52961) using BsmBI restriction sites. For lentivirus production, 2×10^6^ HEK293T cells were transfected with 5 µg lentiCRISPR plasmid (NTC or on target), 1µg pVSV-G (viral envelope) and psPAX2 (2^nd^ generation lentiviral packaging plasmid) using JetPrime (Polyplus, #101000015) according to the manufacturer’s protocol. Six hours after transfection the medium was replaced, incubated for 18 hours and harvested for viral infection of recipient cells. The recipient MDA-MB-468 cells were cultured for 24 hours with lentivirus-containing medium supplemented with 8 µg/ml Polybrene, for an additional 24 hours with fresh medium and selected for 4 days with medium supplemented with 0.75μg/mL puromycin.

After infection, SCD ko pools and clones were cultured in DMEM/F-12 with 10% FBS and 0.8 g/L AlbuMAX II lipid-rich BSA (Thermo Fisher Scientific, # 11021029). After infection, ACSL3 and LRP8 ko pools were cultured in DMEM/F-12 with 10% FBS and 50 nM Na_2_SeO_3_. In maintenance culture, ACSL3 and LRP8 ko clones were grown with 2.5% FBS. sgPSTK, sgSEPHS2 and sgSEPSECS cell lines were cultured in DMEM/F-12 with 2.5% FBS, 0.8 g/l AlbuMAX II lipid-rich BSA and 1µM Ferrostatin-1 (MERCK, # SML0583-5MG) unless otherwise indicated.

Firefly luciferase expression cassette (fluc+) was excised from pGL4.50 vector (Promega, # E1310) using NdeI and BamHI restriction enzymes and cloned into pLenti6NEO vector using the same restriction sites. The integration of the cloned fragment was confirmed by Sanger sequencing. Virus production and infection of MDA-MB-468 target cells was performed as described above for lentiCRISPR vectors. After the infection cells were selected for 7 days with 500 μg/ml G-418S sulphate (Formedium, # G418S). Luciferase activity was checked by supplementing the culture medium with luciferin (100 µg/ml, Abcam, #ab143655) and the bioluminescent signal assessed with a Tecan Spark multiplate reader.

### C11 BODIPY lipid peroxidation assay

5×10^5^ NTC control and SCDko cells were seeded in DMEM/F-12 supplemented with 2.5% FBS in plates of 145mm diameter. 2 days after seeding cells were incubated for 30 min with 1 µM BODIPY 581/591 C11 lipid peroxidation sensor (Thermo Fisher Scientific, #D3861). Cells were then washed with PBS, detached by trypsinisation, pelleted by centrifugation, and resuspended in 400 µL PBS with 1 µg/ml DAPI used to stain dead cells. Peroxidation of the BODIPY probe in live cells was measured by fluorescent-activated flow cytometry (FACS). The fluorescence of the reduced probe was measured at 581/591 nm (excitation/emission, Texas Red filter set) and the oxidised probe at 488/510 nm (FITC filter set). The ratio between the signals at 510 and 591 nm (oxidised/reduced) was used as a readout for lipid peroxidation. 5×10^5^ NTC control and SCDko cells exposed to 100 nM RSL3 for 2 hours were used as positive control for lipid peroxidation.

### Immunoblotting

Cells were lysed in radioimmunoprecipitation assay (RIPA) buffer (Millipore, #20-188). Lysates were incubated in Laemmli buffer (Bio-rad, #1610747) at 95⁰C for 3 min and loaded onto a SDS-polyacrylamide gel (4-12%, Invitrogen NuPAGE, #NP0336BOX). After size separation, proteins were transferred onto nitrocellulose membrane (0.2 µM pore size, Amersham, # 10600001) and membrane was blocked for 1h at room temperature by 5% non-fat dry milk (in tris-buffered saline with 0.01% Tween (TBST)). Membranes were incubated overnight at 4⁰C in a 5% BSA/TBST solution of primary antibody at the following dilutions: vinculin, 1:2000, Merck, # SAB4200080; GPX4, 1:1000, Abcam, # ab125066; ACSL3, 1:1000, Abcam, #ab151959; LRP8, 1:1000, Abcam, # ab108208; SEPHS2, 1:1000, Proteintech, # 14109-1-AP; SCD, 1:1000, Alpha Diagnostics # SCD11-A in Figure 4E and Figure 5G for MDA-MB-468; SCD, 1:1000, Abcam, # ab19862 in Figure 4H and Figure 5G for BT549, CAL120, MDA-MB-231, MCF7. The next day membranes were washed and stained with species-specific near-infrared fluorescent, secondary antibodies (Li-COR) for 1h at room temperature. After additional washing steps, membranes were imaged with Li-Cor Odyssey® DLx imaging system.

**Figure 3:**
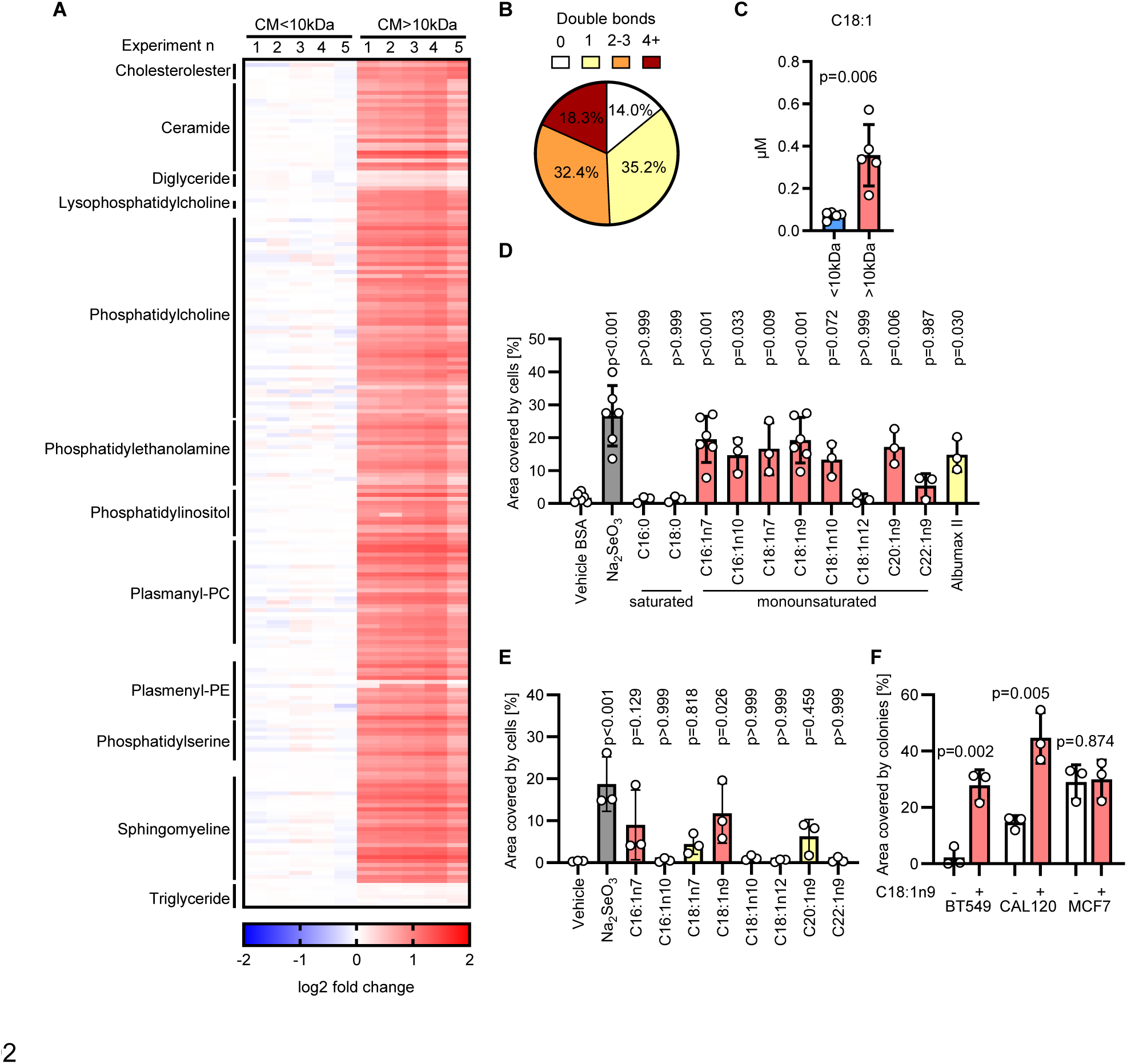
Monounsaturated fatty acids are enriched in the conditioned medium and prevent ferroptosis. A. Heat map of lipids identified in the fractions of condition medium (CM) described in Figure 2B-C. Selected classes of lipids are indicated. n_exp_=5. B. Pie chart depicting the proportion of lipid-bound fatty acids with 0, 1, 2-3, or ≥4 double bonds detected by LC-MS in the CM> 10 kDa fraction . C. Concentration of free C18:1 fatty acid measured in the fractions of conditioned medium described in Figure 2B-C. P value refers to a two-tailed, homoscedastic Student’s *t* tests for unpaired samples. D. Well area covered by colonies formed by MDA-MB-468 cells incubated for 7 days with medium supplemented with 0.015% BSA (vehicle), 0.015% BSA + 50 nM Na_2_SeO_3_, 0.015% BSA + 50 µM of the indicated fatty acids, 1.6g/L lipid-rich BSA (Albumax II). P values refer to a one-way ANOVA test for unpaired samples with Dunnett’s multiple comparisons test. E. Well area covered by colonies formed by MDA-MB-468 cells incubated for 7 days with fresh medium supplemented with vehicle control, 50 nM Na_2_SeO_3_ or 10 µM of the indicated fatty acid. P values refer to a one-way ANOVA test for unpaired samples with Dunnett’s multiple comparisons test. F. Well area covered by colonies formed by the indicated cell lines incubated for 7 days with or without the supplementation of 10 µM oleic acid (C18:1n9). P values refer to a two-tailed, homoscedastic Student’s *t* tests for unpaired samples. C-F. n_exp_=3-5 as indicated by the data points. Bars represent mean ± s.d..

**Figure 4:**
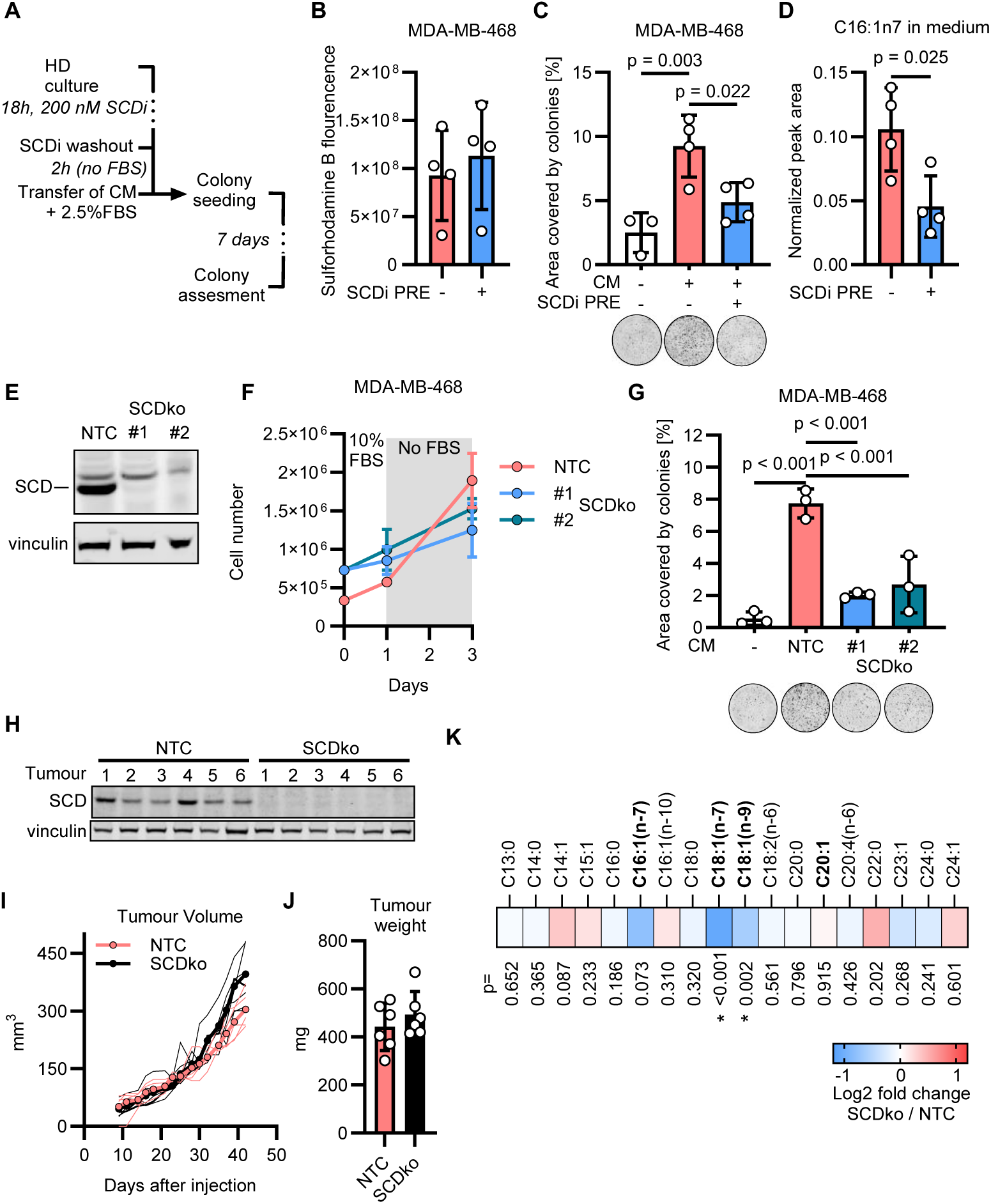
SCD is required for the anti-ferroptotic capacities of conditioned medium. A. Schematic depicting the experimental workflow to condition the medium by MDA-MB-468 at high density (HD) pre-incubated with SCD inhibitor (SCDi, CAY10566). B. Quantification of Sulforhodamine B staining performed on MDA-MB-468 cells immediately after the conditioning of the medium that followed the SCD inhibitor pre-treatment (SCDi PRE) as described in A. C. Well area covered by colonies formed by MDA-MB-468 cells incubated for 7 days with mock medium and medium conditioned for 2 hours by MDA-MB-468 cells without or with SCD inhibitor pre-treatment (SCDi PRE). P values refer to a one-way ANOVA test for unpaired samples with Dunnett’s multiple comparisons test. Representative images of wells with colonies are shown for each experimental condition. D. Quantification of total palmitoleic acid levels (free and lipid-bound C16:1n7) in medium conditioned by MDA-MB-468 cells without or with SCD inhibitor pre-treatment (SCDi PRE). Peak area values are normalized on the signal from internal standard (C17:0). P value refers to a two-tailed, homoscedastic Student’s *t* tests for unpaired samples. E. Immunoblot of SCD and vinculin (loading control) in NTC and two SCDko clones from MDA-MB-468 breast cancer cells. Images representative of 3 independent experiments are shown. F. Number of cells obtained with MDA-MB-468 NTC and SCDko clones cultured as described in the methods section ‘Conditioned medium’. The number of seeded NTC and SCDko cells was adjusted to reach a comparable conditioning of the medium employed for the colony forming assays shown in G. n_exp_=2 G. Well area covered by colonies formed by MDA-MB-468 cells incubated for 7 days with mock medium or medium conditioned by NTC or SCDko MDA-MB-468 clones cultured as described in F and in the methods section ‘Conditioned medium’. P values refer to a one-way ANOVA test for unpaired samples with Dunnett’s multiple comparisons test. Representative images of wells with colonies are shown for each experimental condition. H. Immunoblot of SCD and vinculin (loading control) in NTC and SCDko tumours (n=6) harvested 7 weeks after that 3×10^6^ MDA-MB-468 cells were transplanted unilaterally in the mammary fat pad of NSG female mice. I. Calliper-measured volume of the tumours obtained as described in H. Thinner lines represent the volumes of tumours obtained from each transplantation, thicker lines and symbols represent group means. n_tumour_=6. J. Ex vivo tumour weight of NTC and SCDko tumours described in H. n_tumour_=6. Bars represent mean ± s.d. K. Quantification of total (free and lipid-bound) fatty acid species in interstitial fluid of the tumours described in H. SCD-derived fatty acids are highlighted in bold. Peak area values normalized on the signal from internal standard (C17:0) were used to calculate the Log2 fold change. * indicate significance and p values refer to a two-tailed, homoscedastic Student’s *t* tests for unpaired samples. n_tumour_=6. B-D, G. n_exp_=3-4 as indicated by the data points in each panel. Bars represent mean ± s.d.

**Figure 5:**
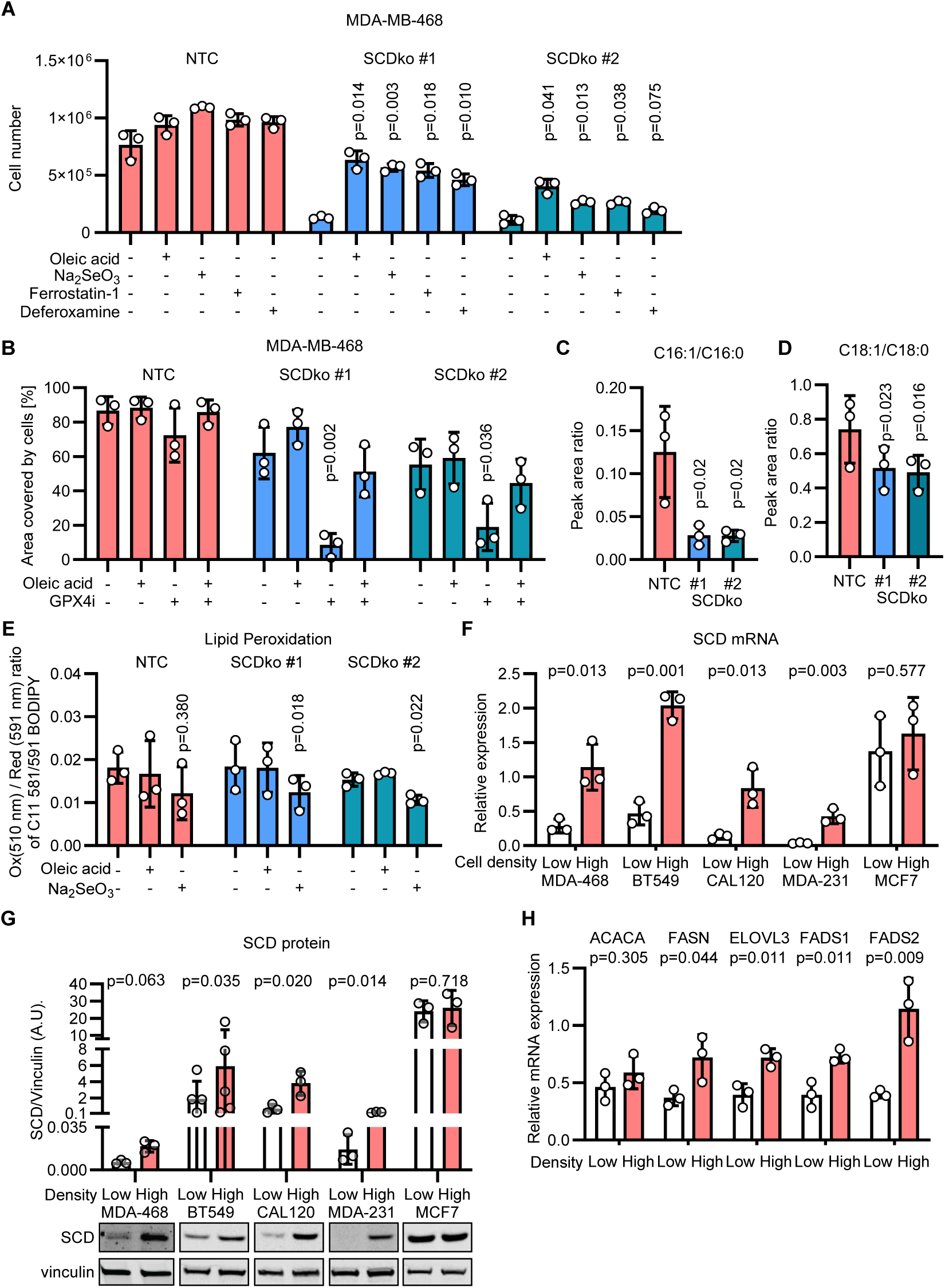
Loss of SCD sensitises cells to ferroptosis. A. Number of NTC and SCDko MDA-MB-468 cells seeded at high density and incubated for 5 days without or with 10 µM oleic acid, 50 nM Na_2_SeO_3_, 2µM Ferrostatin-1 or 2.5 µM Deferoxamine as indicated. P values refer to a two-way ANOVA test for unpaired samples with Dunnett’s multiple comparisons test comparing to the respective unsupplemented controls. B. Well area covered by NTC and SCDko MDA-MB-468 cells seeded at high density and incubated for 5 days without or with 50 nM GPX4 inhibitor (RSL3), 10 µM Oleic acid or their combination (50 nM + 10 µM, respectively). P values refer to a two-way ANOVA test for unpaired samples with Dunnett’s multiple comparisons test comparing to the respective unsupplemented controls. C-D. Peak Area ratio for C16:1 / C16:0 (C) and C18:1 / C18:0 (D) fatty acids in NTC and SCDko MDA-MB-468 cells seeded at high density. P values refer to a one-way repeated measures ANOVA test for paired samples with Dunnett’s multiple comparisons test comparing to the NTC controls. E. Ratio between oxidised (510 nm) and reduced (591 nm) BODIPY 581/591 C11 (lipid peroxidation sensor) in NTC and SCDko MDA-MB-468 cells seeded at high density and incubated for 2 days without or with 10 µM oleic acid or 50 nM Na_2_SeO_3_ as indicated. P values refer to a one-way repeated measures ANOVA test for paired samples with Dunnett’s multiple comparisons test, comparing to the respective unsupplemented controls. F. qPCR quantification of *SCD* mRNA expression in MDA-MB-468, BT549, CAL120, MDA-MB-231 and MCF7 cells seeded at low and high density and cultured for 2 days. The expression is relative to Actin, lamin B1 and TBP mRNA abundance. P values refer to a two-tailed, homoscedastic Student’s *t* tests for unpaired samples. G. Immunoblot analysis of SCD levels in MDA-MB-468 (antibody Alpha Diagnostics, #SCD11-A, 1:1000) and BT549, CAL120, MDA-MB-231, MCF7 cells (antibody Abcam, #ab19862, 1:1000) seeded at low and high density and cultured for 2 days. The lower inset shows representative images of the western blot for SCD and vinculin (loading control) from one of the experiments quantified in the upper panel. P value refers to a two-tailed, homoscedastic Student’s *t* tests for unpaired samples comparing low and high densities. H. qPCR quantification of *ACACA, FASN, ELOVL3, FADS1* and *FADS2* mRNA expression in MDA-MB-468 cells seeded at low and high density and cultured for 2 days. The expression is normalized on the mean mRNA abundance of ACTB, LMNB1 and TBP (**Figure S5 B-D**). P values refer to a two-tailed, homoscedastic Student’s *t* tests for unpaired samples comparing low and high densities. A-H. n_exp_=3-6 as indicated by the data points in each panel. Bars represent mean ± s.d..

### RNA isolation and qRT-PCR analysis

The same number of MDA-MB-468 cells (1.6 ×10^5^) were seeded per well in a 6 well plate or in plates of 145mm diameter for high- and low-density cultures, respectively. For BT549, CAL120, MDA-MB-231 and MCF7 cells, low- and high-density cultures were achieved by seeding 4×10^4^ cells in plates of 145mm diameter and 1.6×10^5^ per well in a 6 well plate, respectively. 2 days after seeding, the cells were washed with ice-cold PBS, scraped off the plate and pelleted at 4⁰C at 10,000g for 30s. RNA was isolated from cell pellets and tumour fragments following the kit manufacturer’s protocol (Qiagen RNeasy, # 74104). 500 ng RNA was used for cDNA synthesis (SuperScript VILO MasterMix, # 11755-050). 5ng cDNA and 8 pmol of each primer were used in each quantitative real-time polymerase chain reaction (Applied Biosystems Fast SYBR Green Master Mix, #4385612). Primer sequences were obtained from primer bank (https://pga.mgh.harvard.edu/primerbank/). A standard curve method with linear regressions R^2^> 0.8 was used to obtain relative quantification of mRNAs expression.

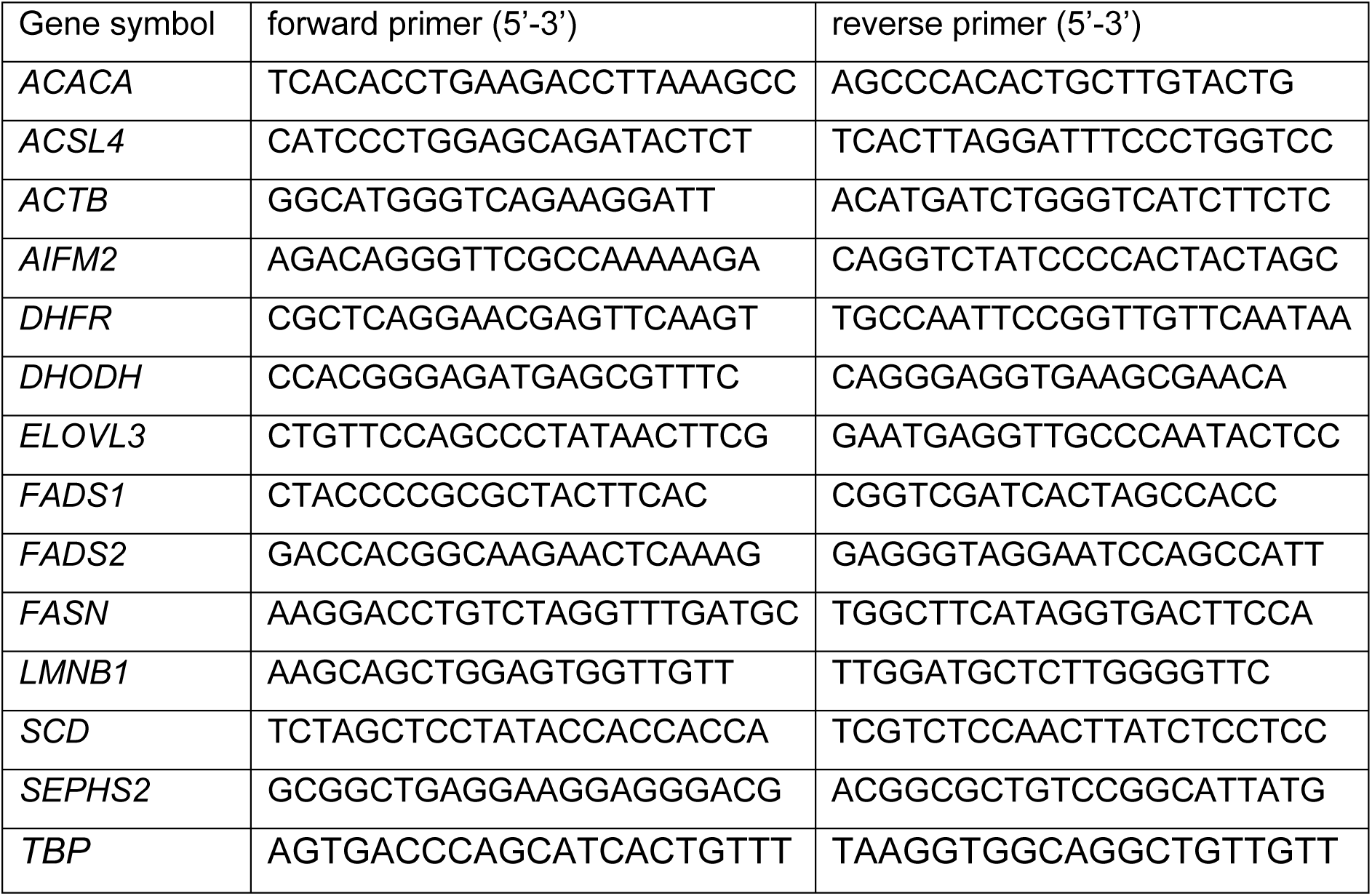

### Xenograft experiments

Animal experiments were performed in accordance with UK Home Office Regulations and Directive 2010/63/EU and subjected to review by the Animal Welfare and Ethical Review Board of the University of Glasgow (Project licence PP6345023 and P38F4A67E). In house-bred or commercially sourced (Charles River) NOD SCID gamma (NSG) mice were housed at temperatures between 19 and 23 ⁰C in ventilated cages with *ad libitum* food and water access and 12 hours light/dark cycles. To minimise pain and distress, Rimadyl was added to the drinking water 24 hours prior to xenotransplantation and removed 3 days post implantation. For experiments with sgSEPHS2 and sgSEPSECS MDA-MB-468 cells, 24 female NSG mice aged between 81-170 days were anesthetized and transplanted into the inguinal mammary fat pad with 50 µl per transplantation of 1:1 PBS:Matrigel solution containing 3×10^6^ luciferase-expressing mycoplasma-negative cells. Groups of eight age-matched animals were randomly assigned to the three experimental groups (NTC, sgSEPHS2 and sgSEPSECS). All the mice were transplanted with cells bilaterally. Tumours were measured by calliper 3 times/week by animal technicians blinded to the scientific outcome. The tumour volume was calculated using the equation [length × width^2^]/2, where width is the smaller of the two dimensions. 38 days post-transplantation mice were culled, and mammary tumours were harvested. Tissues were frozen at -80⁰C or fixed in 10% buffered formalin solution and embedded in paraffin.

To assess the metastatic seeding of breast cancer cells, 2×10^6^ luciferase-expressing mycoplasma-negative MDA-MB-468 cells were passed through a 70 μm strainer, resuspended in 100μL of 4.5% BSA PBS solution (pH 7.4), and injected into the tail vein of 24 female NSG mice aged to 79 days. Mice were randomly assigned to three experimental groups (NTC, sgSEPHS2 and sgSEPSECS) consisting of eight mice per group. Mice were imaged by IVIS bioluminescence at the specified times (indicated in figures) and culled 7-weeks after injection prior to reaching clinical endpoint. Organs were harvested, fixed in 10% buffered formalin solution, and embedded in paraffin for immunohistochemistry.

For experiments with SCDko MDA-MB-468 cells,12 female NSG mice of 113-137 days of age were anesthetized and transplanted unilaterally into the inguinal mammary fat pad with 50 µl per transplantation of 1:1 PBS:Matrigel solution containing 3×10^6^ cells (mycoplasma-negative). Two groups of six age-matched animals were randomly assigned to the NTC and SCDko experimental groups. Tumours were measured by calliper 3 times/week by animal technicians blinded to the scientific outcome. The tumour volume was calculated using the equation [length × width^2^]/2, where width is the smaller of the two dimensions. 43 days post-transplant mice were culled, and tumours were harvested. Tissue fragments were immediately processed for the isolation of interstitial fluid, or frozen on dry ice and stored at -80⁰C or fixed in 10% buffered formalin solution and embedded in paraffin.

### Isolation of interstitial fluid

Tissue freshly isolated from mammary fat pad tumours were cut by scalpel in 4 slices of approximately 2-4 mm thickness and each slice was transferred to an individual cell strainer (Mini Cell Strainer II, Funakoshi, #HT-AMS-04002). The cell strainers were then placed into 1.5ml Eppendorf tubes and centrifuged at 100g for 10 min at 4⁰C to separate the interstitial fluid. The interstitial fluid from 4 slices was merged into one sample representative of one tumour, frozen on dry ice and stored at -80⁰C until lipidomic analysis.

### Immunohistochemistry

All immunohistochemistry (IHC) staining was performed on 4µm formalin fixed paraffin embedded sections (FFPE) heated at 60⁰C for 2 hours.

The following antibodies were used to stain sections with a Leica Bond Rx autostainer: Cas9 (14697, Cell Signaling) and Ku80 (2180, Cell Signaling). All FFPE sections underwent on-board dewaxing (AR9222, Leica) and antigen retrieval using ER2 solution (AR9640, Leica) for 20 min at 95°C. Sections were rinsed with Leica wash buffer (AR9590, Leica) before peroxidase block was performed using an Intense R kit (DS9263, Leica) for 5 min. After rinsing with wash buffer, mouse Ig blocking solution (MKB-2213, Vector Labs) was applied to Cas9 sections for 20 min. Sections were rinsed with wash buffer and then primary antibody applied at an optimal dilution (Cas9, 1:250; Ku80, 1:400) for 30 min. The sections were rinsed with wash buffer and appropriate secondary antibody was applied for 30 min (Cas9, Mouse Envision (Agilent, K4001); Ku80, Rabbit Envision (Agilent, K4003). The sections were rinsed with wash buffer and visualised using DAB with the Intense R kit.

The sections were washed in water and counterstained with haematoxylin z (RBA-4201-00A, CellPath). To complete IHC staining FFPE sections were rinsed in tap water, dehydrated through graded ethanol’s, and placed in xylene. The stained sections were coverslipped in xylene using DPX mountant (SEA-1300-00A, CellPath). Slides were scanned at 20x magnification and analysed with HALO image analysis software (Indica labs).

### Statistical analysis

Independent experimental replicate numbers and statistical tests are described in the figure legends. For all t-tests and ANOVA analyses, GraphPad Prism 9.0 or later versions were used. For the statistical analysis in figure 6F, we used the CGGC permutation test https://bioinf.wehi.edu.au/software/compareCurves/ to assess the significant differences between growth curves.

**Figure 6:**
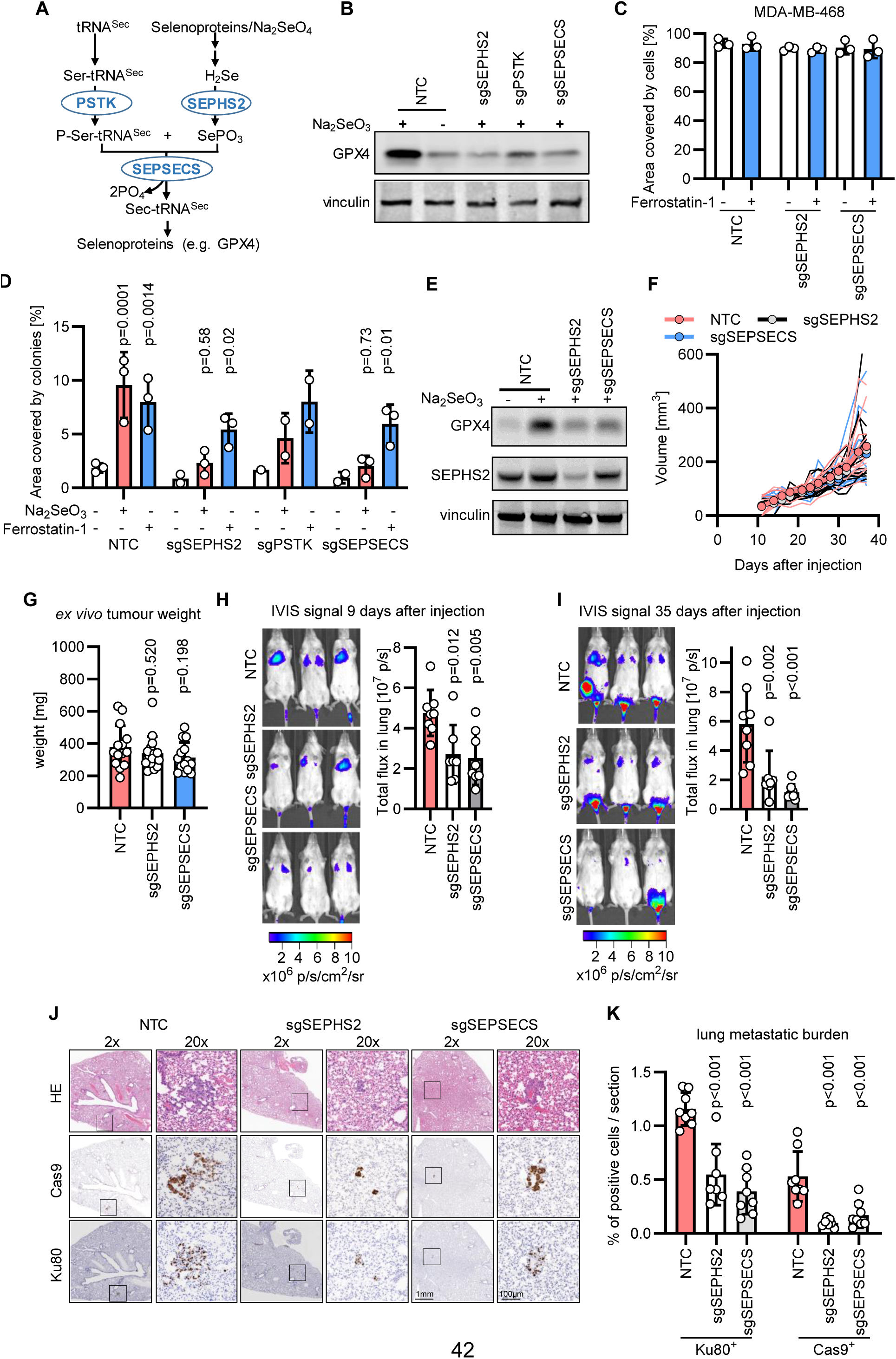
Targeting selenocysteine biosynthesis impairs lung metastasis of TNBC. A. Schematic depicting selenocysteine synthesis on tRNA^sec^. B. Immunoblot images of GPX4 and vinculin (loading control) in control MDA-MB-468 cells (NTC) or in cells depleted of PSTK, SEPHS2 and SEPSECS supplemented with 50 nM Na_2_SeO_3_ as indicated. C. Well area covered by NTC, sgSEPHS2 and sgSEPSECS MDA-MB-468 cells seeded at high density and incubated for 5 days without or with 1 µM Ferrostatin-1. n_exp_=3. Bars represent mean ± s.d.. D. Well area covered by colonies formed by NTC, sgSEPHS2, sgPSTK and sgSEPSECS MDA-MB-468 cells seeded at low density and incubated for 7 days without and with 50 nM Na_2_SeO_3_ or 1 µM Ferrostatin-1. n_exp_=1-3 as indicated by the data points. P values refer to a two-way ANOVA test for unpaired samples with Dunnett’s multiple comparisons test comparing to the respective unsupplemented controls. Bars represent mean ± s.d.. E. Immunoblot images of GPX4, SEPHS2 and vinculin (loading control) in MDA-MB-468 cells (NTC) or in cells depleted of SEPHS2 or SEPSECS supplemented with 50 nM Na_2_SeO_3_ as indicated. Images are representative of 3 independent experiments. F. Calliper-measured volume of tumours in the mammary fat pad of female NSG mice transplanted with NTC, sgSEPHS2 or sgSEPSECS MDA-MB-468 cells (n=8 mice/group). All mice were transplanted bilaterally with 3×10^6^ cells. One transplanted mouse of each NTC and sgSEPSECS group had to be culled before the completion of the experiment for licence reasons due to the location of the tumour. When multiple tumours were formed from a single transplantation of cells their combined volumes were reported as one data point. Thinner lines represent the volumes of tumours obtained from each fat pad transplantation, thicker lines and symbols represent group means (n=14 tumours for NTC, n= 16 for sgSEPHS2, n=14 for sgSEPSECS). A CGGC permutation test (https://bioinf.wehi.edu.au/software/compareCurves/) was used to assess the significant differences between growth curves. Adjusted P value comparing the NTC to sgSEPHS2 or sgSEPSECS obtained with a test with 1000 permutations were 0.66 and 0.67, respectively. G. E*x vivo* weight of resected NTC, sgSEPHS2 or SEPSECS tumours taken 38 days after transplantation. P values refer to a one-way ANOVA test for unpaired samples with Dunnett’s multiple comparisons test comparing to the NTC control. n=14 for NTC or sgSEPSECS tumours and n=16 for sgSEPHS2 as indicated by the data points. H-I. IVIS pictures and quantification of lung metastasis burden 9 days (H) and 35 days (I) after tail vein injection of 2×10^6^ NTC, sgSEPHS2 or sgSEPSECS MDA-MB468 cells. The same mice are shown in H and I. P value refers to a one-way ANOVA test for unpaired samples with Dunnett’s multiple comparisons test comparing to the NTC control. n=7-8 female NSG mice as indicated by data points. One injected mouse of sgSEPHS2 group had to be culled due to husbandry reasons. J. Representative images of staining with haematoxylin and eosin (HE), human Ku80 and Cas9 of lungs of mice described in H-I seven weeks after tail vein injection. Black squares frame the areas magnified at 20x. K. Lung metastasis burden assessed by quantifying the % of human Ku80 and Cas9 positive cells in lungs of mice described in H-I seven weeks after tail vein injection. P values refer to a one-way ANOVA test for unpaired samples with Dunnett’s multiple comparisons test comparing to the NTC control. n=7-8 female NSG mice as indicated by data points.

### Data availability

This study includes no data deposited in external repositories. Source data files that support the findings of this study are stored at the Cancer Research UK Scotland Institute and are available from the corresponding author upon reasonable request. Requests for unique biological materials can be made to the corresponding author.

## Results

### Medium conditioned by breast cancer cells increases colony formation efficiency of triple negative breast cancer (TNBC) cells

We have previously shown that TNBC cells seeded at low density are enabled to form colonies by the sodium selenite present in the physiological cell culture medium Plasmax™ ^23^. To directly visualize if TNBC cells die when seeded at low density we performed live cell imaging on MDA-MB-468 cells (**Figure 1A**). Four days after seeding, the frequency of cell death was significantly decreased if the medium was supplemented with sodium selenite or Ferrostatin-1, demonstrating that TNBC cells seeded at low density die of ferroptosis impairing their colony formation (**Figure 1B**).

However, TNBC cells’ susceptibility to ferroptosis has been shown to be cell density dependent, with cells undergoing ferroptosis selectively when seeded at low density ^23,33,34^. To dissect the mechanism underpinning the dependency of ferroptosis on culture density, we tested whether the ferroptotsis-resistance observed in selenium-restricted cultures at high density (**Figure S1A**) was transferrable with the conditioned medium. The medium conditioned for 48 hours by high-density cultures of MDA-MB-468 cells, was employed to seed cells at low density. Similarly to the anti-ferroptotic agents, sodium selenite and Ferrostatin-1, the conditioned medium conferred the capacity to survive ferroptosis and form colonies to MDA-MB-468 cells (**Figure 1C** and **S1B**). The medium conditioned by high-density cultures of human BT549, CAL-120, MDA-MB-231 and murine EO771 TNBC cells also promoted colony formation in the respective cell lines seeded at low density (**Figure 1D**). However, this effect was not observed with the luminal A, ER-positive MCF7 breast cancer cells, for which the number of colonies was not altered by the conditioned medium (**Figure 1D**). Moreover, the medium conditioned by BT549, CAL-120 and MCF7 cells stimulated MDA-MB-468 colony-forming capacity, demonstrating that the anti-ferroptotic effect of conditioned medium is preserved across cell lines (**Figure 1E**).

This observation prompted us to test if conditioned medium from untransformed cells could also support clonal growth of cancer cells. Cancer-associated fibroblasts (CAFs) are known to support the proliferation of cancer cells *in vivo* ^27,35^. Conditioned medium from CAFs increased the colony formation capacity of MDA-MB-468 comparably to the anti-ferroptotic agent sodium selenite (**Figure 1F**). Conversely, neither the media conditioned by human normal fibroblasts from the mammary gland (MF), or from the dermis (DF), significantly increased the colony-forming capacity of MDA-MB-468 cells (**Figure 1F**). These data suggest that the medium conditioned by transformed or cancer-reprogrammed cells supports clonal growth of cancer cells by preventing ferroptosis.

### Breast cancer cells cultured at high density produce and secrete an anti-ferroptotic factor

To test whether cells cultured at high density condition the medium by depleting a ferroptosis inducing molecule (e.g. iron) or produce a factor that prevents ferroptosis, we collected medium conditioned by high density MDA-MB-468 cells and diluted with fresh medium (1:3). The diluted conditioned medium enabled colony formation of MDA-MB-468 cells comparably to medium supplemented with the anti-ferroptotic agent Ferrostatin-1 (**Figure 2A**). This factor is also produced and secreted by TNBC cells cultured in a sodium selenite-free version of the physiological culture medium Plasmax™ ^23^ (**Figure 2A**). These results suggest that breast cancer cells produce an anti-ferroptotic factor that compensates for the selenium deficiency. To test whether the conditioned medium regulates the expression of known ferroptotic mediators in the recipient cells we measured the mRNA levels of AIFM2, ACSL4, DHFR, DHODH, and GPX4 in cells cultured at low density and exposed to the condition medium. However, the expression pattern of these genes did not explain the anti-ferroptotic activity of the conditioned medium (**Figure S2A-F**), suggesting that the secreted factor does not activate a transcriptional anti-ferroptotic response in the recipient cells.

To identify this factor, we first fractionated the conditioned medium with a size exclusion column with a 10 kDa cut-off (**Figure 2B**). The flow-through fraction of the condition medium depleted of molecules > 10 kDa did not significantly increase the colony forming capacity of MDA-MB-468 cells compared to unconditioned medium. On the contrary the concentrated fraction of the conditioned medium enriched in molecules > 10 kDa significantly boosted the colony formation when used to supplement the fresh medium (**Figure 2C**). Moreover, its activity was retained when incubated for 15 min at 95°C (**Figure 2D**). To further investigate the chemical properties of this factor we applied a Bligh and Dyer lipid extraction to the > 10kDa fraction of the condition medium. The extracted lipidic fraction used to supplement the medium of the MDA-MB-468 cells increased their colony forming capacity comparably to sodium selenite supplementation (**Figure 2E**). Altogether these results suggest that the active anti-ferroptotic factor produced by TNBC cells at high density is bound to a dispensable carrier > 10 kDa and is a heat-resistant lipid sufficient to confer anti-ferroptotic effects to the conditioned medium. Consistently with this hypothesis, the deletion of the long-chain-fatty-acid-CoA ligase 3 (ACSL3), an enzyme that plays a key role in lipid biosynthesis, ablated the protective effect of the conditioned medium (**Figure 2F-G**), strengthening the hypothesis that the secreted pro-survival factor is of lipidic nature.

### Monounsaturated fatty acids prevent ferroptosis of TNBC cells

To gain an insight into the lipidic composition of the conditioned medium, we performed a lipidomic analysis on its size-separated fractions. Overall, we could detect 213 lipids from 12 classes, and all of them were more abundant in the > 10 kDa fraction (**Figure 3A**). Notably, 86% of the acyl chains of the detectable lipids had one (35%) or more double bonds (**Figure 3B**). In addition to the lipidic classes reported in Figure 3a, we could detect free C18:1 monounsaturated fatty acid (MUFA), that was significantly more abundant in the > 10 kDa fraction of the conditioned medium (**Figure 3C**).

Based on this evidence, we tested if individual fatty acid (FA) supplementation influenced the capacity of MDA-MB-468 cells to form colonies. At 50 µM, the saturated FA 16:0 and 18:0 did not have a significant effect on colony formation, while out of the 8 MUFAs tested, 7 significantly increased the colony forming capacity (**Figure 3D and S3B**). Of the 7 MUFAs with colony stimulating activity, 4 (C16:1n7, C18:1n7, C18:1n9 and C20:1n9) retained their activity at a lower concentration (10 µM, **Figure 3E and S3C**). In addition, 10 µM C18:1n9 (oleic acid) increased the colony formation capacity in BT549 and CAL120 TNBC cells but not in MCF7 cells, thereby phenocopying the effects of conditioned medium across these breast cancer cell lines (**Figure 3F and S3D**).

To profile the effects of oleic acid and conditioned medium on the lipidome of recipient cells we performed lipidomic analysis on MDA-MB-468 cells cultured at low cell density and exposed to those agents. The results show that 350 out of the 790 lipidic species identified were consistently and coherently regulated by the oleic acid and the conditioned medium compared to cells exposed to mock medium (**Figure S3A**). Triglycerides emerged as the class of lipids whose abundance was consistently increased by both oleic acid and conditioned medium. Conversely, polyunsaturated phosphatidylcholines were downregulated in cells cultured with conditioned medium or oleic acid. These results show that the lipidome changes elicited by oleic acid and the condition medium largely overlap, suggesting a common mode of action for these anti-ferroptotic agents.

### SCD activity is required to condition the medium with an anti-ferroptotic factor

Notably, all the MUFAs stimulating the clonogenic growth at 10 µM (**Figure 3E**) were products of the stearoyl-CoA desaturase (SCD), therefore we hypothesized that the presence of the anti-ferroptotic factor in the conditioned medium could be SCD-mediated. To test this hypothesis, we compared the colony-stimulating capacities of media conditioned by MDA-MB-468 cells pre-treated with the SCD inhibitor CAY10566 or vehicle control (**Figure 4A**). SCD inhibition had no significant effects on the viability of high-density cultures employed to condition the medium (**Figure 4B**). Nevertheless, the medium conditioned by SCD-inhibited cells was significantly less effective in stimulating colony formation compared to medium conditioned by vehicle-treated cells (**Figure 4C**). Consistently with this observation, the levels of C16:1n7 (palmitoleic acid), a product of SCD, were significantly lower in the medium conditioned by SCD-pre-inhibited cells (**Figure 4D**) while other fatty acids were not significantly affected (**Figure S4A**).

To further validate the essential role of SCD to produce the anti-ferroptotic factor secreted in the medium, we generated two SCD knockout (ko) clones derived from MDA-MB-468 cell lines (**Figure 4E**). Constitutive *SCD* deletion (SCDko) impaired cell proliferation (**Figure 4F**), therefore the number of cells seeded was adjusted to obtain comparable levels of medium-conditioning in 48 hours (**Figure 4F**). Compared to medium conditioned by NTC cells, the media from SCDko cells was significantly less potent in stimulating colony formation (**Figure 4G**). Moreover, media conditioned by SCDko cells was less effective than medium from NTC cells in supporting colony formation of cells that lack the selenoprotein P (SELENOP) receptor, LRP8 (**Figure S4B-C**). These results indicate that SCD activity in TNBC cells is required to produce an anti-ferroptotic factor transferrable with the conditioned medium, whose action is not dependent on the uptake of a major selenium carrier, SELENOP.

To test if the SCD-dependent secretion of lipids occurs in the tumour microenvironment *in vivo*, SCDwt and SCDko MDA-MB-468 cells were orthotopically transplanted in the mammary fat pad of mice and the interstitial fluid was collected from the respective tumours. The deletion of SCD (**Figure 4H**) did not affect tumour growth (**Figure 4I-J)**, but it selectively decreased the levels of the SCD-derived MUFAs oleic, palmitoleic and vaccenic acids in the tumour interstitial fluid (**Figure 4K)**. These results show that breast cancer cells release SCD-produced MUFAs in the extracellular tumour microenvironment where, similarly to what we observed in high cell density cultures, MUFAs and selenium-dependent pathways have overlapping anti-ferroptotic functions.

### Cell density-dependent loss of SCD expression sensitizes TNBC cells to ferroptosis

Having demonstrated the anti-ferroptotic role of SCD-derived lipids secreted in the conditioned medium, we assessed whether SCD-deletion could sensitize to ferroptosis TNBC cells cultured at high density upon selenium restriction (i.e. 2.5% FBS). The number of SCDko cells cultured at high density was partially rescued by the individual supplementation of oleic acid, sodium selenite, ferrostatin-1 and deferoxamine (**Figure 5A**) demonstrating that SCDko promotes ferroptosis upon selenium restriction. In addition, the survival of the SCDko cells, but not that of NTC control cells, was significantly impaired by the treatment with 50 nM of GPX4 inhibitor RSL3 and rescued by oleic acid supplementation (**Figure 5B and S5A**), demonstrating that the SCD-dependent production of MUFAs such as palmitoleic and oleic acids (**Figure 5C-D**) becomes essential for cell survival when the selenium-dependent anti-ferroptotic function is hindered. Unexpectedly, the levels of oxidized C11 BODIPY, a probe for lipid peroxidation, were not significantly affected by *SCD* deletion (**Figure 5E**) suggesting that SCD-derived fatty acids desensitize cells from ferroptosis acting downstream of lipid peroxides.

To find a mechanistic rationale for the cell density-dependent susceptibility to ferroptosis observed in TNBC cells (i.e. maximal at low density), we assessed the levels of SCD mRNA and protein expression and found that it was proportional to cell density in all four TNBC lines tested (**Figure 5F-G**). Consistently with the unresponsiveness of luminal A MCF7 cells to the conditioned medium (**Figure 1B**), these cells did not modulate *SCD* expression depending on cell density (**Figure 5F-G**). Furthermore, the mRNA expression of key enzymes for fatty acid synthesis, elongation, and desaturation (FASN, ELOVL3, FADS1 and FADS2) was also proportional to TNBC cell density (**Figure 5H**).

These data show that TNBC cells have an impaired fatty acid synthesis and desaturation capacity at low cell density, thereby increasing their vulnerability to ferroptosis mediated by selenium starvation.

### Inhibition of selenocysteine synthesis induces ferroptosis and impairs lung metastatic colonization

GPX4 is a potent anti-ferroptotic enzyme whose catalytic activity requires selenocysteine in its active site. The amino acid selenocysteine is only synthesised on its tRNA and cannot be re-cycled as such for the synthesis of selenoproteins (**Figure 6A**). This makes selenocysteine synthesis a limiting step to counteract ferroptosis and it could constitute a novel therapeutic target for ferroptosis-primed TNBC cells. On these bases, we designed guide RNAs (gRNAs) against the three genes encoding the enzymes required for the synthesis of selenocysteinilated tRNA^sec^ (PSTK, SEPHS2 and SEPSECS, **Figure 6A**). The interference with the expression of each of the enzymes resulted in low GPX4 levels in MDA-MB-468 cells, comparable to those observed upon selenium starvation (**Figure 6B**). The loss of SEPHS2 or SEPSECS expression as well as Ferrostain-1 supplementation had no significant effects on the viability of cells seeded at high density, demonstrating that the anti-ferroptotic function of selenoproteins is redundant under these conditions (**Figure 6C and S6A**). On the contrary, at low cell density survival depends on ferroptosis inhibition, achieved by either selenium or ferroststatin-1 supplementation (see NTC in **Figure 6D and S6B**). In these ferroptosis-priming conditions, the interference with SEPSECS, SEPHS2 and PSTK expression ablates the protective effect of selenium, while Ferrostatin-1 retains its anti-ferroptotic effect. These results indicate that selenocysteine synthesis inhibition triggers ferroptosis selectively in TNBC cells cultured at low density (**Figure 6D and S6B**).

Next, we orthotopically transplanted NTC control, sgSEPHS2 and sgSEPSECS MDA-MB-468 cell pools into the mouse mammary fat pad to investigate the effect of selenocysteine synthesis on tumour growth (**Figure 6 E-F**). The growth rate of sgSEPHS2 and sgSEPSECS tumours was not different from that of NTC controls, an observation confirmed ex vivo by tumour weights (**Figure 6F-G**). However, compared to NTC controls the lower levels of SEPHS2 and GPX4 expression observed pre-implantation (**Figure 6E**) were restored in sgSEPHS2 or sgSEPSECS tumours at endpoint (**Figure S6D-E**), suggesting a counterselection of the interfered cells.

To assess whether the colonization of distant organs by TNBC cells is affected by selenocysteine synthesis inhibition, we injected luciferase-expressing NTC, sgSEPHS2 and sgSEPSECS MDA-MB-468 cells into the tail vein of female NSG mice. One hour after tail vein injection, the luciferase signal was detectable in the lungs of all mice, and it was comparable between experimental groups (**Figure S6C**). However, nine days after injection, the luciferase signal was significantly lower (∼2-fold) in animals injected with sgSEPHS2 and sgSEPSECS compared to NTC cells (**Figure 6H**). This difference was further exacerbated 35 days after injection (∼3-5 fold, **Figure 6I**). To validate the luciferase-based results, we stained the lungs of the tail vein-injected animals with antibodies against Cas9 and human Ku80 to visualize individual metastatic cells (**Figure 6J**). We found two to three times more Ku80-positive cells in lungs of mice injected with NTC cells (∼1.15% of all cells in the lung) compared to sgSEPHS2(∼0.55%) or sgSEPSECS cells (∼0.4%, **Figure 6K**). The magnitude of these differences was enhanced if Cas9-postive cells were counted in the lungs of the same mice (NTC cells ∼0.55%, sgSEPHS2 ∼0.1%, and sgSEPSECS ∼0.2%). Overall, these *in vivo* results demonstrate that the expression of SEPHS2 and SEPSECS is dispensable for primary breast tumour growth but support the capacity of breast cancer cells to survive the bloodstream and colonize the lungs, recapitulating the conditional essentiality of selenocysteine synthesis for clonogenic growth observed *in vitro*.

## Discussion

Understanding the regulation of ferroptotic cell death will provide new avenues to treat cardiovascular disease, neurodegenerative diseases, and cancer ^36,37^. In the recent years, several redundant anti-ferroptosis pathways have been described ^3–5,7^. However, it remains to be elucidated which of those pathways are conserved in specific tissues and cell types. We and others have previously showed that selenium deprivation and uptake inhibition sensitizes to ferroptosis selectively when cells are cultured at low density ^23,24,34^. Here we demonstrate that in triple-negative breast cancer (TNBC) cells, the seeding density positively regulates the expression of genes involved in fatty acid synthesis and desaturation (i.e. SCD). Intriguingly, we found that medium conditioned by cancer cells and cancer associated fibroblasts cultured at high density has a pro-survival effect on TNBC cells seeded at low density for colony forming assays (**Figure 1**). Specifically, we showed that the pro-clonogenic activity of the conditioned medium is thermostable and is retained in its lipidic fraction (**Figure 2E**). Moreover, the supplementation of specific monounsaturated fatty acids (MUFAs) was effective in preventing ferroptosis occurring during colony formation (**Figure 3D**). Indeed, SCD-derived MUFAs are the most potent in rescuing ferroptosis when supplemented to TNBC cells at low density. Consistently, the loss of SCD activity in TNBC cells cultured at high density sensitises to ferroptosis (**Figure 5A-C**). These results suggest that diets low in SCD-derived MUFAs might enhance the efficacy of pro-ferroptotic agents in TNBC ^15^. In line with this hypothesis, Dierge et al. showed that oleate-rich and PUFA-rich diets have opposite effects on the ferroptosis susceptibility of acidotic cancer cells, and Ubellacker et al. described the anti-ferroptotic effects of oleic acid in melanoma ^16,38^. We showed that unlike normal cells, high density cultures of breast cancer cells or cancer-associated fibroblasts secrete lipids containing monounsaturated fatty acid in the extracellular environment in quantities that prevent ferroptosis when supplemented to colony-forming cells (**Figure 1 and 2E**). Recently, it has been shown that primary breast tumours can secrete a factor that increases palmitate production in the lung, conditioning the metastatic niche to favour cancer cell growth ^39^. Furthermore, it has been shown that the pro-metastatic effect of high fat diets relies on CD36-dependent MUFA uptake ^40^. Our results show that mammary tumours, where cells are at high density, secrete MUFA-containing lipids in the interstitial fluid altering the lipid composition of tumour microenvironment (**Figure 4K** and Synopsis). On these bases, we hypothesized that selenocysteine biosynthesis could be a novel target to kill ferroptosis-primed triple-negative breast cancer (TNBC) cells that have left the MUFA-rich environment of the primary tumour. Indeed, the interference with SEPHS2 or SEPSECS, enzymes of the selenocysteine biosynthesis pathway (**Figure 6A**), strongly impairs the lung seeding of cancer cells injected into the bloodstream (**Figure 6H-K**). In culture, we show that each of the three enzymes synthesizing the Sec-tRNA^sec^ (PSTK, SEPHS2 and SEPSECS) as well as selenium supplementation are all required to prevent ferroptosis in cells at low density (**Figure 6**). Carlisle et al. have shown that high SEPHS2 expression correlates with poor survival in patients with breast carcinoma and that its loss induces glioblastoma cell death by the accumulation of toxic H_2_Se upon supraphysiological supplementation of sodium selenite ^20^. Our data show that loss of SEPHS2 in TNBC cells phenocopies the cytotoxic effects of selenium deprivation and does not require the accumulation of toxic selenium metabolites to prime cells to ferroptosis (**Figure 6C and D**).

Breast cancer tissue shows increased levels of protein-bound selenium compared to healthy tissue surrounding the tumour ^41^ suggesting that unrestricted selenium availability is promoting tumour cells survival. Indeed, GPX4 inhibition in combination with immunotherapy showed promising results in murine models of TNBC ^42^. While selenium-deprived diets have not been tested in breast cancer models, Eagle et al. showed a therapeutic effect of selenium deprivation in leukaemia models, while Alborzinia et al. inhibited cellular selenoprotein P uptake to decrease the expression of selenoproteins and induce ferroptosis in neuroblastoma^22,24^.

On the contrary, selenium-supplemented diets have been assessed with clinical trials (Select trial, NCT00006392) as preventive interventions for breast and prostate cancer with the rationale to boost the antioxidant capacity of the host. However, whether diet intervention can lead to selenium concentrations that limit selenocysteine synthesis has yet to be established. Moreover, the pharmacologic inhibition of Sec-tRNA^sec^ synthesis might have a more precise effect on the selenoprotein production in cancer cells that enter the blood circulation and depend on the antioxidant function of selenoproteins for their survival. Interestingly, PSTK shows low similarity to other eukaryotic kinases thereby favouring the design of specific small molecule inhibitors ^43^.

Taken together, our *in vitro* data show that SCD expression increases with cell density in TNBC cells and mechanistically, the loss of the pro-survival effect of SCD in cells at low density enhances their dependency on the antioxidant action of selenoproteins to survive ferroptosis and form colonies. This principle translates *in vivo* where we show that TNBC cells injected in the bloodstream metastasize less efficiently to the lung when selenoprotein synthesis is impaired by the deletion of SEPHS2 and SEPSECS, two key enzymes in the Sec-tRNA^sec^ synthesis. In line with the working model proposed the deletion of SEPHS2 and SEPSECS does not impact the proliferation of TNBC cultured at high density or grown as orthotopic mammary tumours, suggesting that that inhibition of the Sec-tRNA^sec^ could be a valid therapeutic target to eradicate metastatic TNBC cells primed to ferroptosis by an imbalance in fatty acid saturation.

## Acknowledgements

The authors thank the Core Services and Advanced Technologies at the Cancer Research UK Scotland Institute, and particularly the Metabolomics, Biological Services Unit, Histology Service, Molecular Technologies and Advanced Imaging Resource. The authors acknowledge Catherine Winchester (CRUK Scotland Institute) for critically reviewing this manuscript and all the members of the Oncometabolism lab for the constructive discussions. This work was funded by Cancer Research UK core funding awarded to the CRUK Scotland Institute (grant number A31287), Stand Up to Cancer campaign for CRUK awarded to S.Z. (grant number A29800), Breast Cancer Now awarded to S.Z. (grant number 2018NovPR102), Cancer Research UK core funding awarded to K.B. (grant number A29799) and Cancer Research UK core funding awarded to S.T. (grant number A23982).

## Author contributions

The study was conceptualized by T.A. and S.T.; methods were developed by E.S., G.R.B., C.N., V.H.V, D.S.; experiments were performed by T.A., E.S., R.D., J.A., L.C.A.G., B.A.S.; data were graphically visualized by T.A. and S.T.; the study was supervised by L.M. S.Z., K.B., D.S., S.T.; the manuscript was written by T.A. and S.T with reviewing and editing by all authors; funding was acquired by S.Z., K.B.,S.T..

## Competing interests

ST is the inventor of Plasmax^TM^ cell culture medium. All other authors declare no competing interests.

---

**Figure S1:**
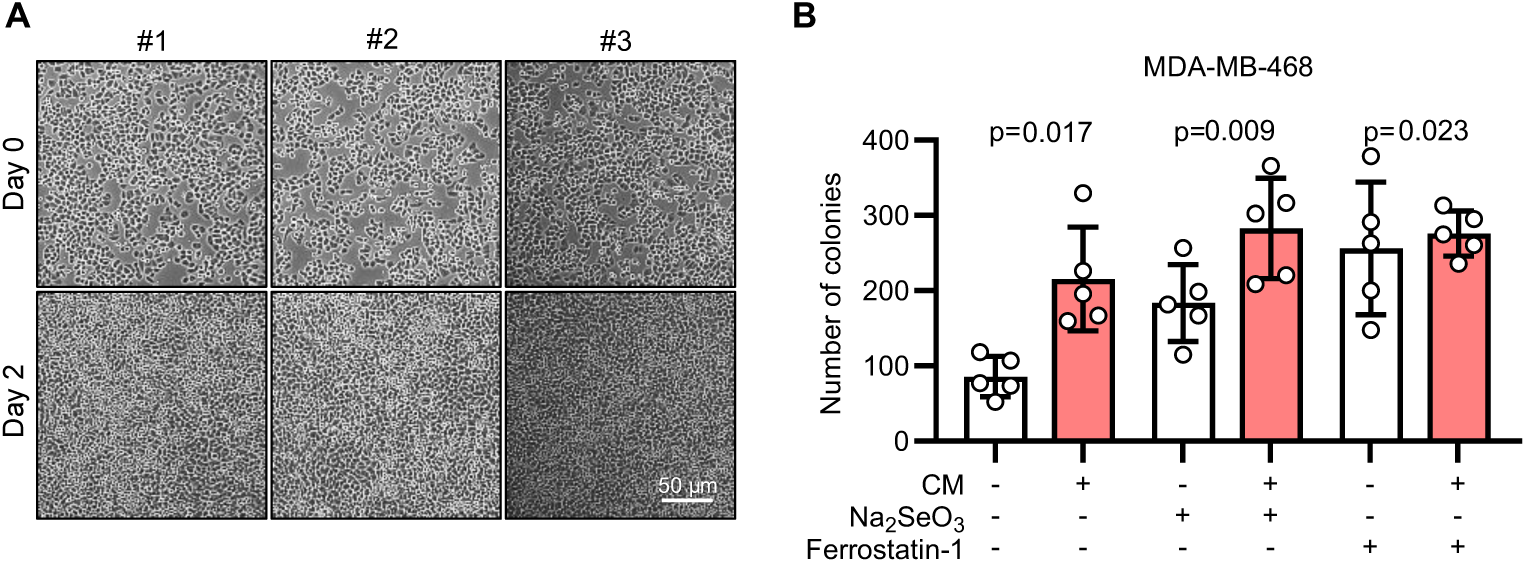
Medium conditioned by breast cancer cells and cancer-associated fibroblasts enhances the clonogenicity of triple-negative breast cancer cells. A. Images of high-density cultures of MDA-MB-468 cells at the start (day 0) and end (day 2) of the medium conditioning period. Representative fields of view from 3 independent experiments (#1-3) are shown. B. Quantification of the number of colonies obtained from the assays shown in Figure 1C. P values refer to a two-way ANOVA test for unpaired samples with Dunnett’s multiple comparisons test

**Figure S2:**
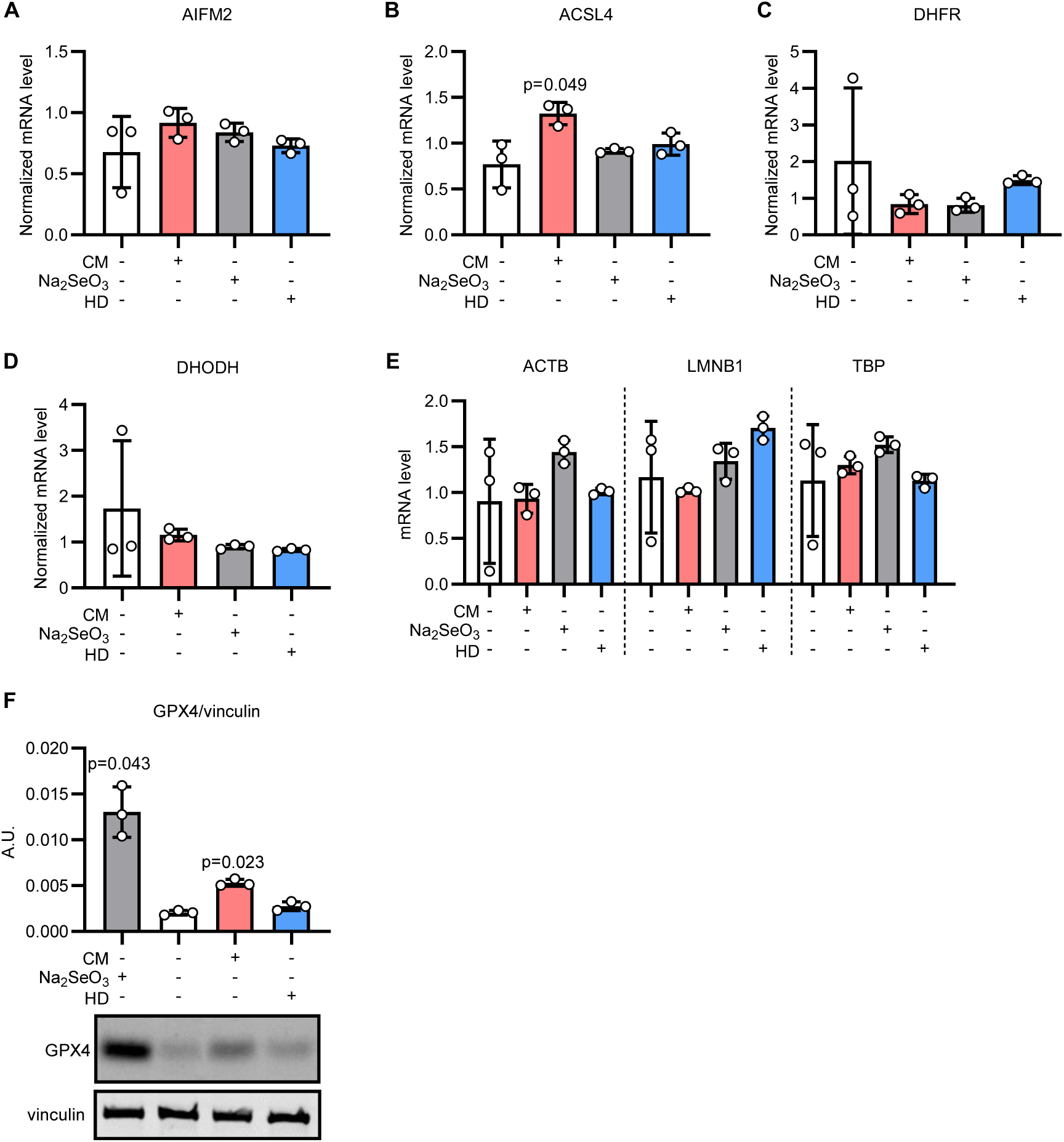
Breast cancer cells produce an anti-ferroptosis molecule at high density. A-D. qPCR quantification of *AIFM2* (A)*, ACSL4* (B)*, DHFR* (C), and *DHODH* (D) mRNA expression in MDA-MB-468 cells seeded at low density in mock medium and supplemented with 50 nM selenite or conditioned medium (CM) for 2 days as indicated. The gene expression was also assessed in cells seeded at high density (HD) grown for 2 days in mock medium. The mRNA level is normalized to the mean mRNA abundance of the 3 housekeeping genes (*ACTB, LMNB1,* and *TBP*) shown in E. P values refer to a one-way ANOVA for paired samples with Dunnett’s multiple comparisons test. n_exp_=3. Bars represent mean ± s.d. E. qPCR quantification of *ACTB, LMNB1,* and *TBP* mRNA expression in MDA-MB-468 cells seeded and treated as described in A-D. F. Immunoblot analysis of GPX4 and vinculin (loading control) in MDA-MB-468 cells MDA-MB-468 cells seeded and treated as described in A-D. A.U.: arbitrary unit. Representative images of GPX4 and vinculin (loading control) from one of the 3 experiments quantified in the upper graph. P values refer to a one-way ANOVA for paired samples with Dunnett’s multiple comparisons test. Bars represent mean ± s.d.

**Figure S3:**
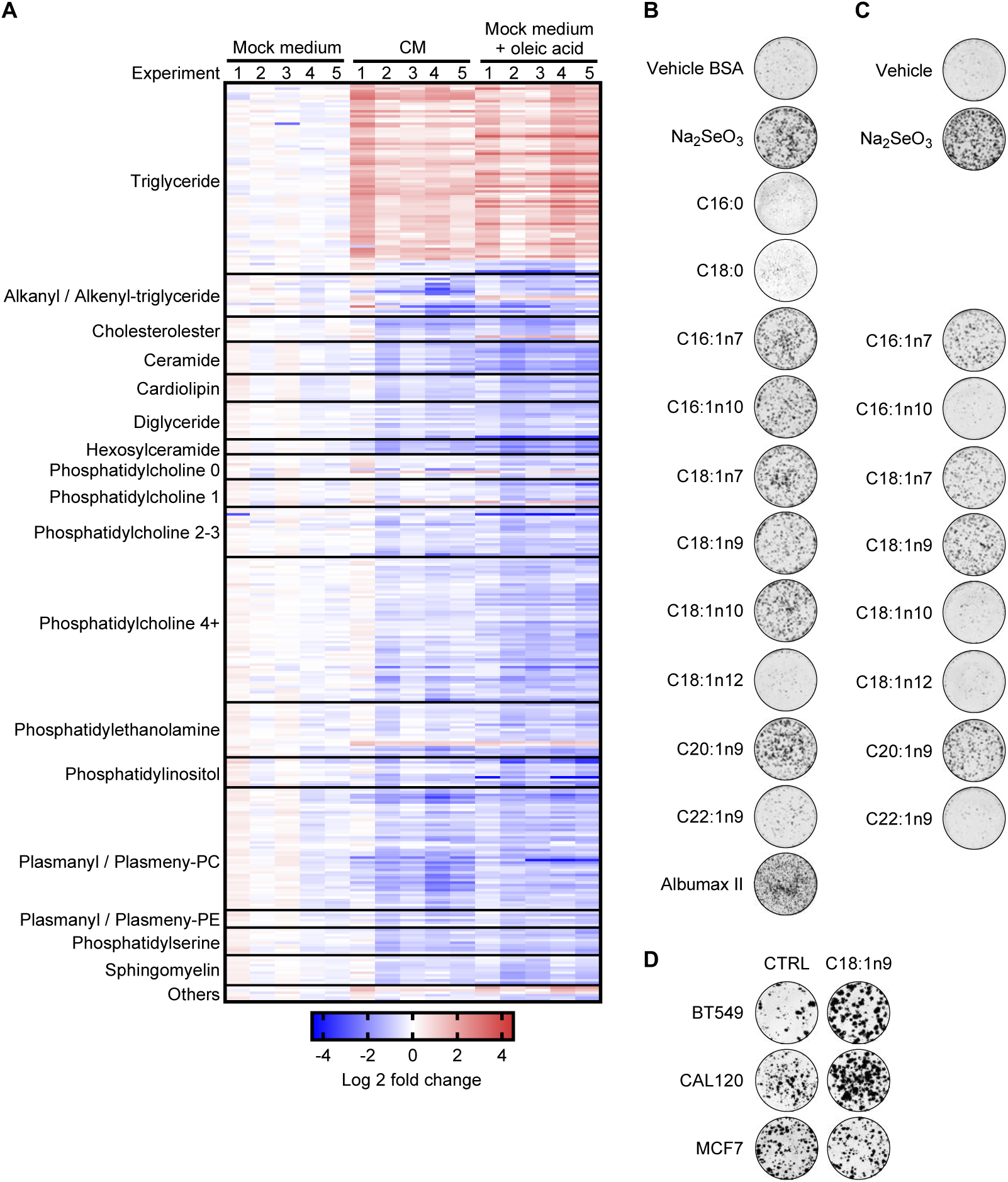
Monounsaturated fatty acids are enriched in the conditioned medium and prevent ferroptosis. A. Heatmap of lipids regulated in MDA-MB-468 cells cultured at low density with mock medium, conditioned medium (CM) or mock medium with 10 µM oleic acid. Ferrostatin-1 was supplemented at 2 µM in all conditions. The lipids identified as significantly regulated with a two-tailed, homoscedastic Student’s *t* tests for unpaired samples in the comparison between conditioned medium and mock medium are reported and selected classes of lipids are indicated. For the phosphatidylcholine class the number of double bonds is also reported (0, 1, 2-3, 4+). The Log2 fold change refers to the comparison with mock medium supplemented cells. n_exp_=5. B. Representative images for the colony forming assays displayed in Figure 3D. C. Representative images for the colony forming assays displayed in Figure 3E. D. Representative images for colony forming assays displayed in Figure 3F.

**Figure S4:**
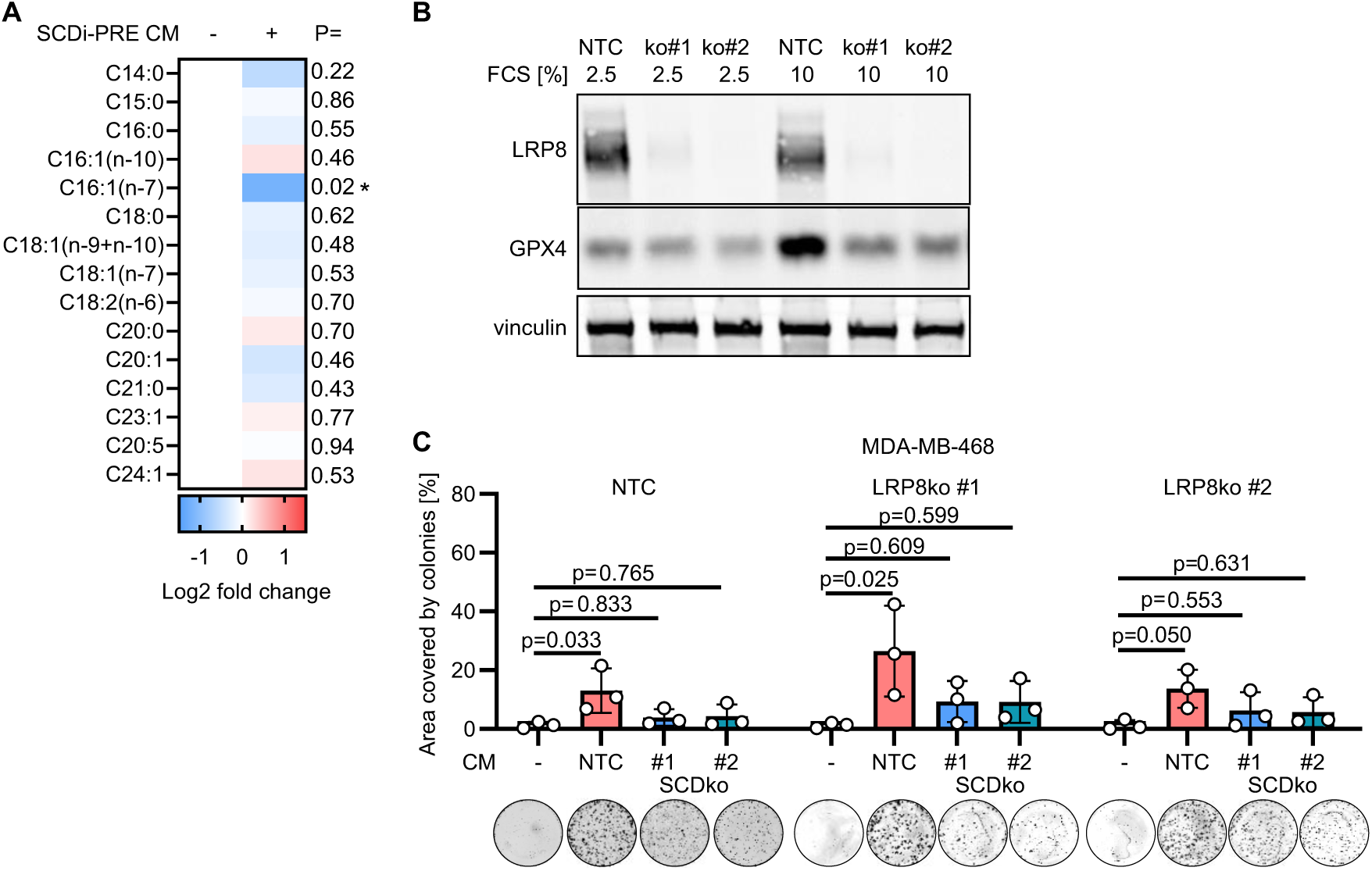
SCD is required for the anti-ferroptotic capacities of conditioned medium. A. Quantification of total (free and lipid-bound) fatty acid species in medium conditioned by MDA-MB-468 cells without or with SCD inhibitor pre-treatment (SCDi PRE). Peak area values normalized on the signal from internal standard (C17:0) were used to calculate the Log2 fold change. P value refers to a two-tailed, homoscedastic Student’s *t* tests for unpaired samples. These data complement Figure 4D. n_exp_=4. B. Immunoblot of LRP8, GPX4, and vinculin (loading control) in NTC and LRP8ko clones (#1-2) derived from MDA-MB-468 breast cancer cells. C. Well area covered by colonies formed by MDA-MB-468 NTC control cells and LRP8ko clones incubated for 7 days with mock medium or medium conditioned by NTC or SCDko MDA-MB-468 clones cultured as shown in Figure 4F. P values refer to a one-way ANOVA test for unpaired samples with Dunnett’s multiple comparisons test. n_exp_=4. Bars represent mean ± s.d.. Representative images of wells with colonies are shown for each experimental condition.

**Figure S5:**
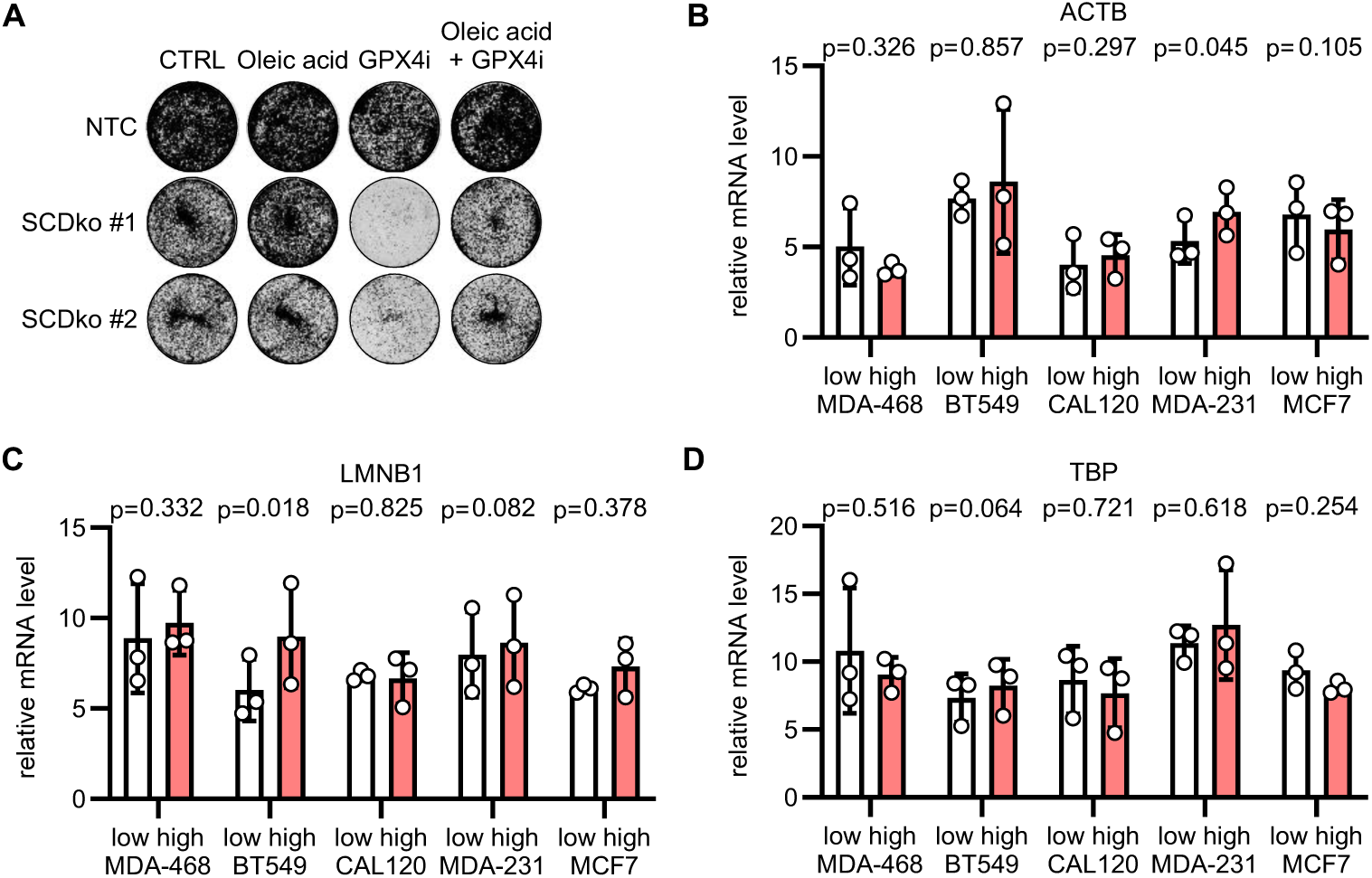
Loss of SCD sensitises cells to ferroptosis. A. Representative images of the colony forming assays shown in Figure 5B. B-D. qPCR quantification of ACTB (B), LMNB1 (C), and TBP (D) mRNA expression in MDA MB-468, BT549, CAL120, MDA-MB-231 and MCF7 cells seeded at low or high density and cultured for 2 days. The mean values for the three genes were used to normalize the expression of the genes shown in Figure 5 H. P value refers to a two-tailed, homoscedastic Student’s *t* tests for paired samples comparing low and high densities. Bars represent mean

**Figure S6:**
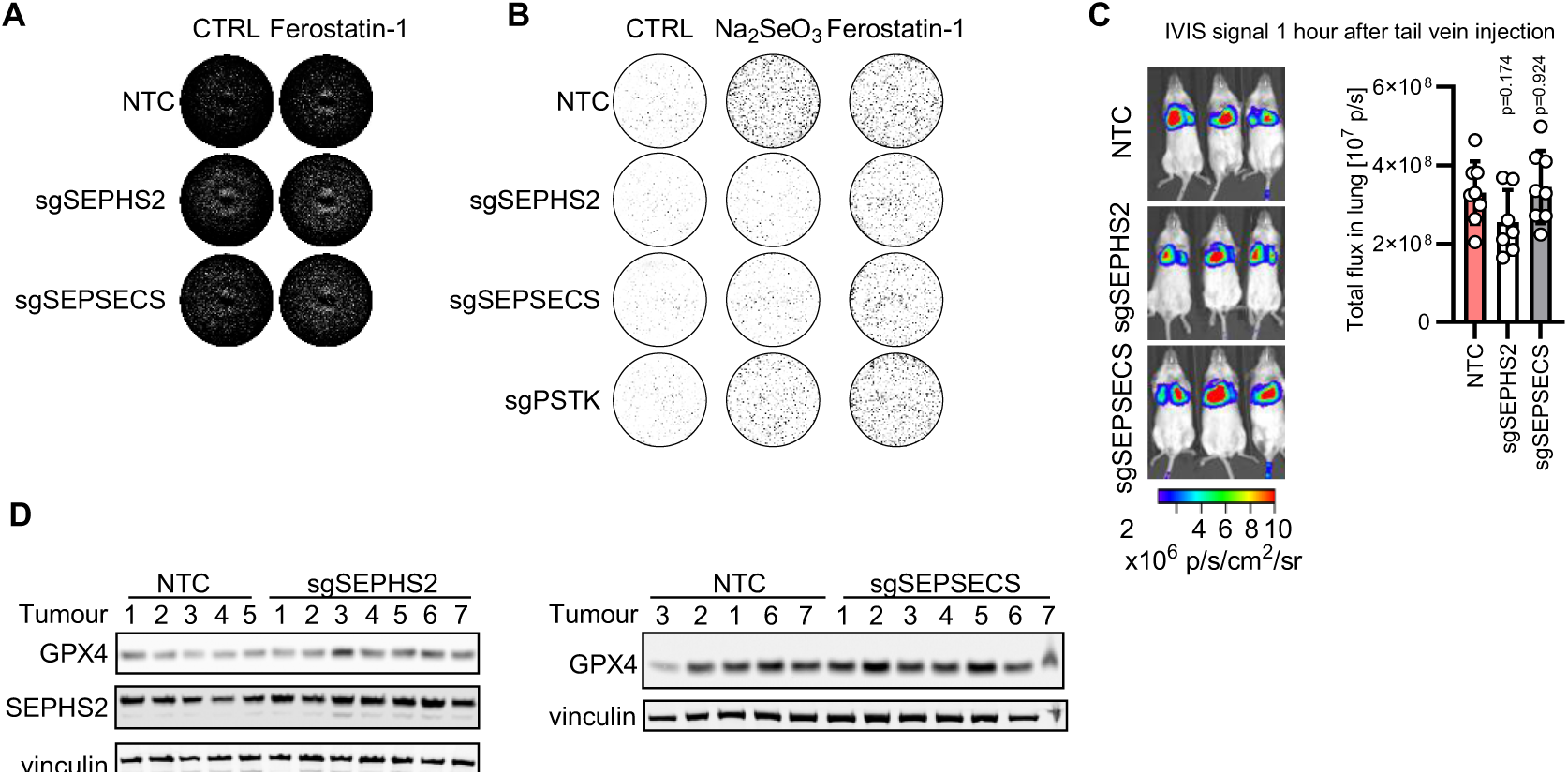
Targeting selenocysteine biosynthesis impairs lung metastasis of TNBC. A. Representative images of the well area covered by cells at the end of the assays shown in Figure 6C. B. Representative images of the colony forming assays shown in Figure 6D. C. IVIS pictures and quantification of lung metastasis burden 1 hour after tail vein injection of 2×10^6^ NTC, sgSEPHS2 or sgSEPSECS MDA-MB468 cells. The same mice are shown in Figure 6 H and I. P value refers to a one-way ANOVA test for unpaired samples with Dunnett’s multiple comparisons test comparing to the NTC control. n=7-8 female NSG mice as indicated by data points. One injected mouse of sgSEPHS2 group had to be culled due to husbandry reasons. D. Immunoblot for GPX4, SEPHS2 and vinculin (loading control) in mammary tumours sampled 38 days after the transplantation of NTC, sgSEPHS2, or sgSEPSECS MDA-MB-468 cells. For each experimental group the lysates from 7 tumours were loaded as indicated.

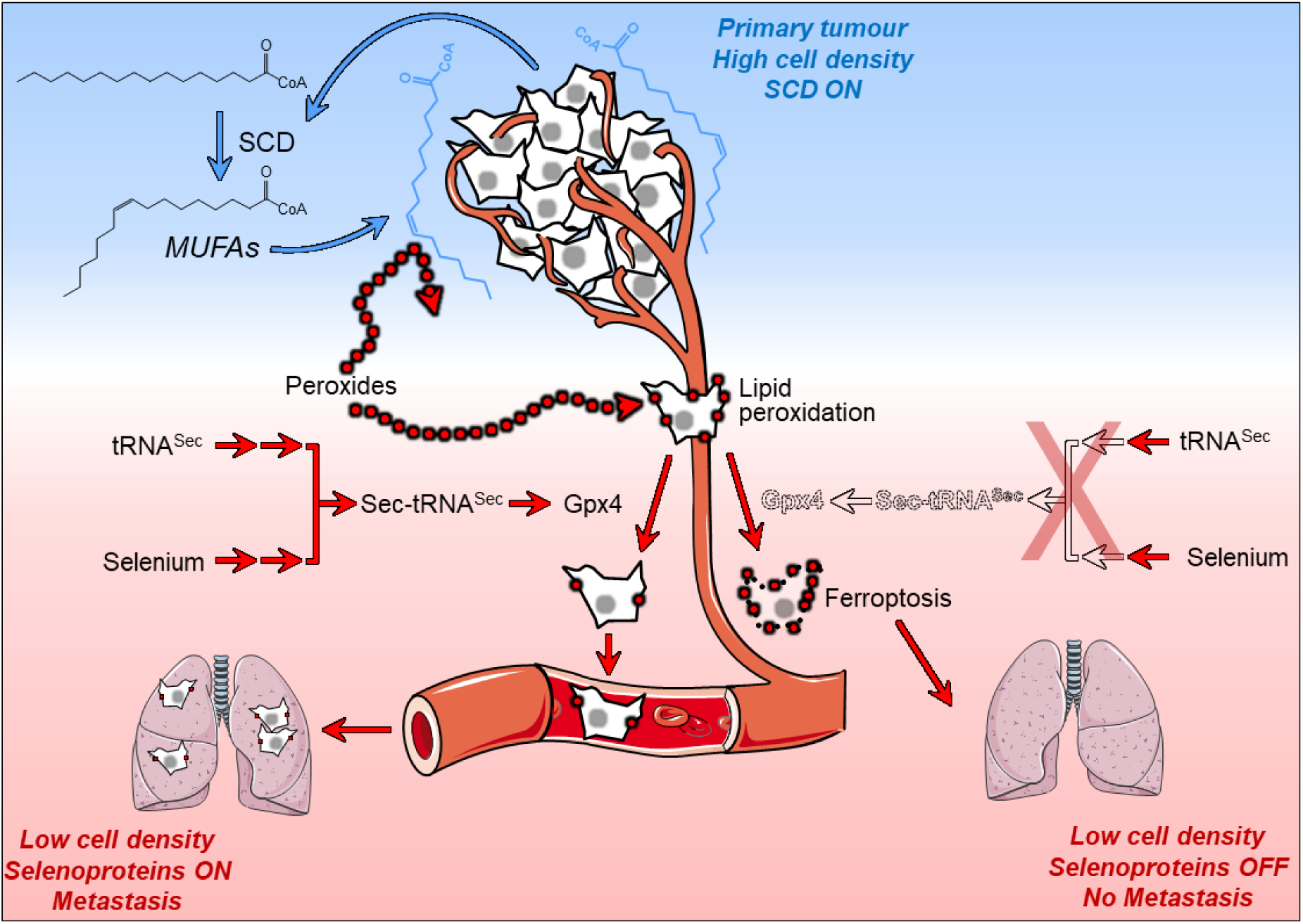
**Synopsis.** Triple negative breast cancer (TNBC) cells grown at high density and in cell-dense mammary tumours produce MUFAs *via* Stearoyl-CoA Desaturase (SCD) and secrete lipid-bound MUFAs that protects them from pro-ferroptotic lipid peroxidation. Conversely, TNBC cells grown at low density or metastasizing in the bloodstream downregulate SCD expression resulting in MUFAs deficiency. This metabolic shift renders metastatic cells dependent on the anti-ferroptotic action of selenoproteins, in particular Glutathione Peroxidase 4 (GPX4), exposing a novel conditional vulnerability tied to the inhibition of Sec-tRNA^sec^ biosynthesis. Indeed, in preclinical models targeting the enzymes of the Sec-tRNAsec biosynthesis effectively impede TNBC lung metastasis.

## Notes

### Competing Interest Statement

ST is the inventor of PlasmaxTM cell culture medium. All other authors declare no competing interests.

### Summary of Updates

We added in live cell imaging data to show that cells die of ferroptosis at low density. (Figure 1A,B). We investigated effects of conditioned medium on LRP8ko and ACSL3ko cells (Figure 2 and S4). We added new in vivo experiments that show that TNBC cells secrete MUFAs to the interstitial fluid (Figure 4). We repeated in vivo experiments with selenocysteine biosynthesis inhibition and added a new target (SEPSECS) to these experiments (Figure 6).

